# How the dual PDZ domain from Postsynaptic density protein 95 clusters ion channels and receptors

**DOI:** 10.1101/775726

**Authors:** Nazahiyah Ahmad Rodzli, Michael Lockhart-Cairns, Colin W. Levy, John Chipperfield, Louise Bird, Clair Baldock, Stephen M. Prince

## Abstract

PSD-95 is a member of Membrane Associated Guanylate Kinase class of proteins which form scaffolding interactions with partner proteins including ion and receptor channels. PSD-95 is directly implicated in modulating the electrical responses of excitable cells. The first two PSD-95/Disks Large/Zona Occludens domains of PSD-95 have been shown to be the key component in the formation of channel clusters. We report crystal structures of the dual domain in both in apo and ligand-bound form; thermodynamic analysis of ligand association and Small Angle X-ray Scattering of the dual domain in the absence and presence of ligands. These experiments reveal that the ligated double domain forms a scaffold in the complete sense of the word. The concentration of the components in this study is comparable to those found in compartments of excitable cells such as the postsynaptic density and juxta-paranodes of Ranvier. The properties of the dual domain explain the basis of the scaffolding function of PSD-95, and provide a more detailed understanding of the integration of key components of neuronal specializations involved in nervous signal transmission.

## Introduction

The organization of ion channels or ionotrophic receptors at high-densities at specialized locations in the cell membrane is fundamental to the function of neurones. These locations include the synapse, the axonal hillock and axonal locations such as nodes of Ranvier in myelinated neurones. The Disks Large homologue 4 protein commonly known as SAP-90 or PSD-95 is abundant in the Post Synaptic Density (PSD) (Li et al, 2004), and in rat has also been found at axonal juxta-paranodes (Ogawa et al, 2010) adjacent to nodes of Ranvier. The PSD is a high staining cytoplasmic layer localized at the surface of the synaptic membrane (Palay, 1956). The area ascribed to the PSD is of the order of 0.05µm^2^ and extends some 35-50nm into the cytoplasm (Harris & Weinberg, 2012). The PSD has a laminar structure with N-methyl D-aspartate (NMDA) selective glutamate receptors in the membrane closely associated with PSD-95 (Valtschanoff & Weinberg, 2001). Knock down studies have shown that the PSD-95 protein plays an important role in the integrity of synapses by anchoring receptors (Chen et al, 2015). The PSD-95 protein has been studied in relation to a number of disorders of the central nervous system including both acute and chronic conditions (Gardoni et al, 2009), which reflects the importance of the protein in neuronal function.

Aspects of the structural role of the PSD-95 protein in the coordination of channels and receptors have been revealed by numerous experiments: Studies in mouse have shown that the PSD-95 protein organizes ionotrophic receptors and ion channels by forming super-complexes on the mega-Dalton scale (Frank et al, 2017). Imaging studies have shown that PSD-95 regulates the organization of α-amino-3-hydroxy-5-methyl-4-isoxazolepropionic acid (AMPA) selective glutamate receptors in the post-synaptic membrane via the formation of domains on the nanometre scale (Nair et al, 2013). Using tomographic studies in combination with antibody-labelling separations of PSD-95 molecules in postsynaptic isolates of ~13 nm have been observed (Chen et al, 2008). *In vitro* studies have shown the formation of complexes on the association of the cytoplasmic domain of an inwardly rectifying Kir2.1 potassium channel and PSD-95 (Fomina et al, 2011).

PSD-95 is the best-characterized member of the Membrane Associated Guanylate Kinase (MAGUK) family (Gomperts, 1996; Won et al, 2017) which also includes PSD-93, SAP-97 and SAP-102. MAGUK family proteins contain five linked domains and both PSD-95 and PSD-93 are localised to the membrane. A schematic diagram of PSD-95 is shown in Figure 1(a). Experiments in rat have shown that PSD-95 localization is via palmitoylation of Cys residues at the amino-terminus (El-Husseini et al, 2000a). The canonical human isoform of PSD-95 comprises 724 amino acids (<u>Uniprot</u> P78352). The domains of PSD-95 are distributed within the primary sequence interspersed with linker peptides of varying lengths (Figure 1(a)). From the N-terminus a 64 residue linker is followed by a PSD-95/Disks Large/Zona Occludens protein (PDZ) domain (PDZ1) a short linker (8 residues) and then a second PDZ domain (PDZ2), a second stretch of 66 amino acids links to a third PDZ domain (PDZ3) followed by a 34 residue linker and closely associated SRC (kinase) Homology 3 (SH3) and Guanylate Kinase (GK) domain.

**Figure 1.**
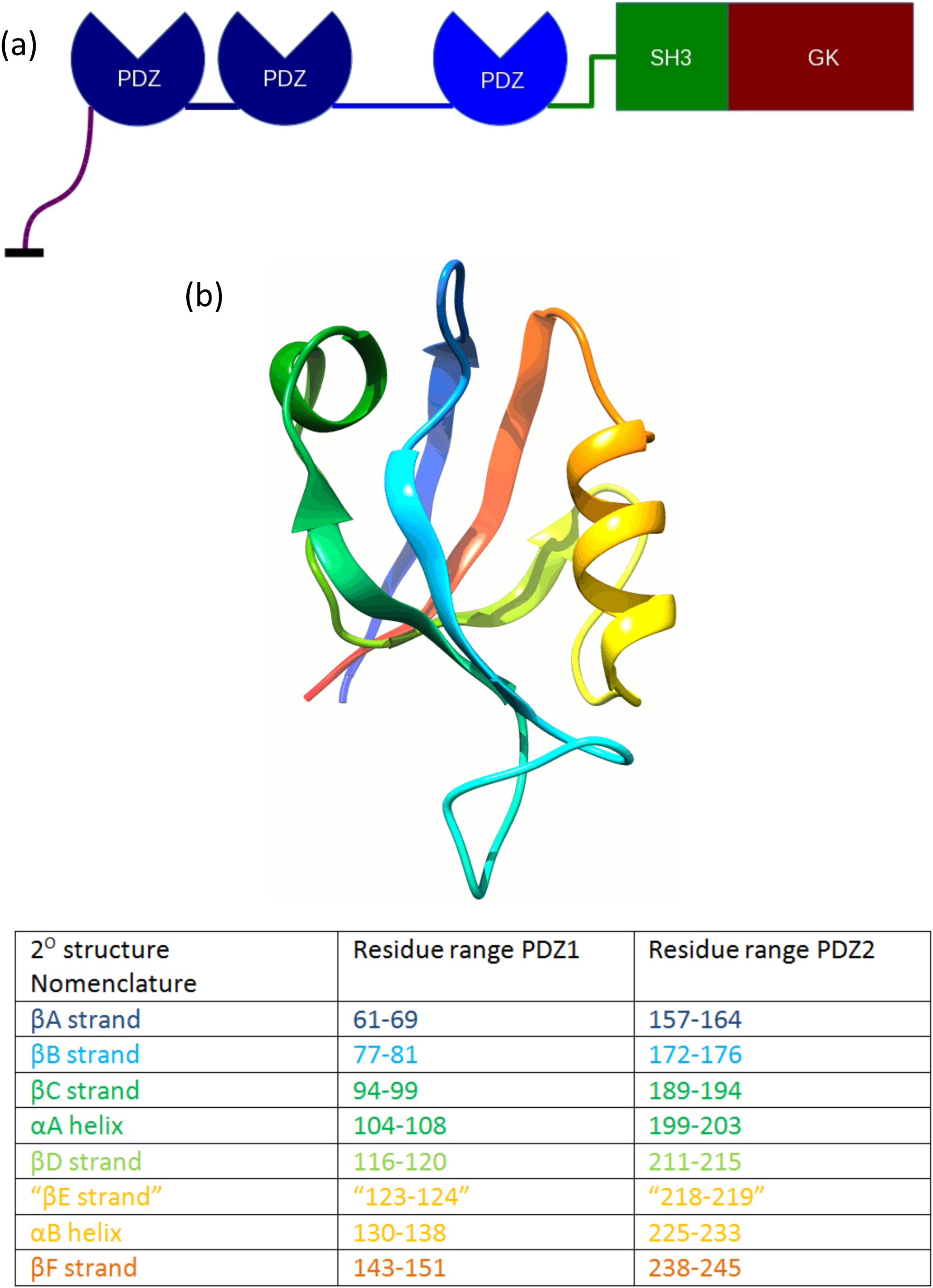
PSD-95 and PDZ domains. A schematic of the overall structure of PSD-95: Inter-domain linkers are shown as lines, PDZ domains are shown as segmented circles. The residue order is indicated by colouring Violet-Red (N-C terminus). (b) A cartoon rendering of a PDZ domain, derived from chain A of PDB entry 3rl7 (Zhang et al, 2011), coloured according to increasing sequence number (N-terminus, blue; C-terminus red). The colour key indicates the secondary structure definitions (Kim & Sheng, 2004) referred to in the accompanying table (the βE strand is enclosed by inverted commas as it is not assigned for this structure). Secondary structure elements are shown in a ribbon representation, and the sequence order of β-strands indicated by an arrowhead. The cartoon diagram was produced by the UCSF Chimera program (Pettersen et al, 2004).

PDZ domains consist of around 90 amino acids and many have a characteristic -Gly-Leu-Gly-Phe-sequence and are sometimes named GLGF domains for this sequence. PDZ domains are abundant in the human genome and are present in over 400 proteins, often with multiple copies (Kim & Sheng, 2004). They are also found in other organisms in some having the role of presenting peptides to proteases (Inagaki et al, 1996). PDZ domains have been classified according to the properties of their cognate ligands (Songyang et al, 1997; Stricker et al, 1997). The PDZ domains of PSD-95 are all class I PDZ domains (Songyang et al, 1997) where the ligand is the Carboxyl-terminus (C-terminus) of the partner protein. The type I PDZ ligand has a consensus sequence of the form –X_-3_–(Ser/Thr/Cys)_-2_-X_-1_-Φ_0_ where X is any residue and the C-terminal residue Φ_0_ has an aliphatic side chain.

A large number of structures of PDZ domains have been determined using Nuclear Magnetic Resonance or X-ray crystallography. A schematic diagram of the domain is shown in Figure 1(b), the binding cleft for the peptide ligand lies between the βB strand and the long αB alpha helix. The GLGF signature sequence of the PDZ domain is found at the apex of the binding cleft with GLG in the βA-βB loop. For type I PDZ domains GLGF interacts with the C-terminus of the partner sequence (Doyle et al, 1996).

A number of ionotrophic synaptic receptors and Potassium ion channels possess PDZ binding motifs at their C-termini, and these have been shown to interact with MAGUKs including PSD-95 (Gomperts, 1996). Recent studies have shown that component proteins of the synapse are able to self-assemble into aggregates and the PSD-95 protein is a component of all of these self-assembling systems (Zeng et al, 2018; Zeng et al, 2016). An earlier study dissected the roles of segments of PSD-95 in clustering Shaker Kv1.4 channels in transfected COS-7 cells (Hsueh et al, 1997). This work found that the N-terminal linker region is essential for clustering and that clustering is observed even for truncated N-terminal constructs, for example a construct containing only the N-terminal linker and PDZ1-2 of PSD-95 mediated clustering with wild-type efficiency. Mutation of Cys residues at residues 3 and 5 of an N-terminal-linker-PDZ1-2 construct was found to abolish co-immunoprecipitation of the truncated protein with full-length PSD-95. This resulted in a hypothesis that inter-molecular disulphide linkage of residues in the N-terminal region could be a mechanism for clustering by PSD-95.

A structure of the PDZ1-2 dual domain in complex with a peptide derived from the C-terminus of the cypin protein (sequence QVVPFSSSV) was obtained via NMR (Wang et al, 2009), (PDB ID 2ka9). The ensemble of structures deposited in this study show variation in the relative orientations and separation of domains in PDZ1-2. Two crystal structures of the PDZ1-2 fragment of PSD-95 have also been solved. A crystal structure of human apo PDZ1-2 from PSD-95 with 4 copies of the dual domain in the asymmetric unit is available at a resolution of 3.4 Å (Bach et al, 2012) (PDB ID 3zrt). The overall conformation of all four copies of 3zrt:PDZ1-2 is essentially identical although part of PDZ1 in one copy of 3zrt:PDZ1-2 is not resolved in the crystal structure. A crystal structure of Rat PDZ1-2 from PSD-95 with two copies of the dual domain in the asymmetric unit is available at a resolution of 2.05Å (Sainlos et al, 2011) (PDB ID 3gsl). A ligating sequence (ETMA) derived from the C-terminus of the ionotrophic glutamate receptor Glur6 is fused to the C-terminus of the 3gsl:PDZ1-2 expressed protein. This -ETMA ligand sequence is observed to associate exclusively to PDZ1 domains of 3gsl:PDZ1-2 in the crystal structure. The gross conformation of the PDZ1-2 domains in 3zrt:PDZ1-2 and 3gsl:PDZ1-2 differ by a rotation along the axis of the 8-residue inter-domain linker. The inter-domain rotation is larger for the 3zrt:PDZ1-2 with respect to that between the two copies of PDZ1-2 in 3gsl:PDZ1-2.

The interaction of full-length PSD-95 and the tetrameric cytoplasmic domain of the inwardly rectifying potassium channel (Kir2.1) leads to the formation of extended molecular complexes as seen in earlier work at low resolution (Fomina et al, 2011). The aim of this study was to increase our understanding of the molecular details of clustering by studying the essential interacting components of this complex, namely the PDZ1-2 fragment pf PSD-95 and the C-terminal peptide sequence of Kir2.1, at high resolution.

## Results

### X-ray crystallography

The detailed interaction of a compatible sequence with both PDZ domains of PDZ1-2 is only resolved using a high-resolution structural technique, hence both PDZ1-2 alone and PDZ1-2 plus a ligand peptide were studied using X-ray crystallography. A sequence of RRESEI corresponding to the last 6 residues of Kir2.1 was used. The ESEI sequence comprises a type I PDZ domain interaction motif and the preceding RR residues were included to ensure that a peptide amino terminus does not interfere with ligand association with the PDZ cleft. These RR residues have also been implicated in receptor trafficking (Standley et al, 2000). The RR residues ensure overall charge neutrality and may enhance peptide solubility. Crystals of un-ligated PDZ1-2 (apo:PDZ1-2) were obtained initially and these were used to micro-seed drops of PDZ1-2 with ligand. Crystal structures in the un-ligated (apo:PDZ1-2) and ligated (RRESEI:PDZ1-2) state were obtained at resolutions of 2.0 Å (R_free_ 26.0%) and 2.1 Å (R_free_ 23.8%) respectively (see Supplementary Table S1). The crystal structures obtained are in the same tetragonal Spacegroup with a very similar unit cell and the same gross PDZ1-2 double domain conformation is present in both apo:PDZ1-2 and RRESEI:PDZ1-2 (Figure 2(a)). The PDZ1-2 complex has a direct intra-molecular contact between PDZ1 and PDZ2 formed by interactions between the βB-βC loop of PDZ2 and the αA helix of PDZ1. A hydrogen bond is formed between the main chain amide O of Ala 106 and the side chain amide NH_2_ of Gln 181 (Nε2-O distance of 3.0Å), and there is a direct interaction between Pro 101 and Pro 184. The αA(PDZ1):βB-βC(PDZ2) interaction results in a more compact conformation of PDZ1-2 in the structures reported here compared to those found in 3zrt:PDZ1-2 and 3gsl:PDZ1-2.

**Figure 2.**
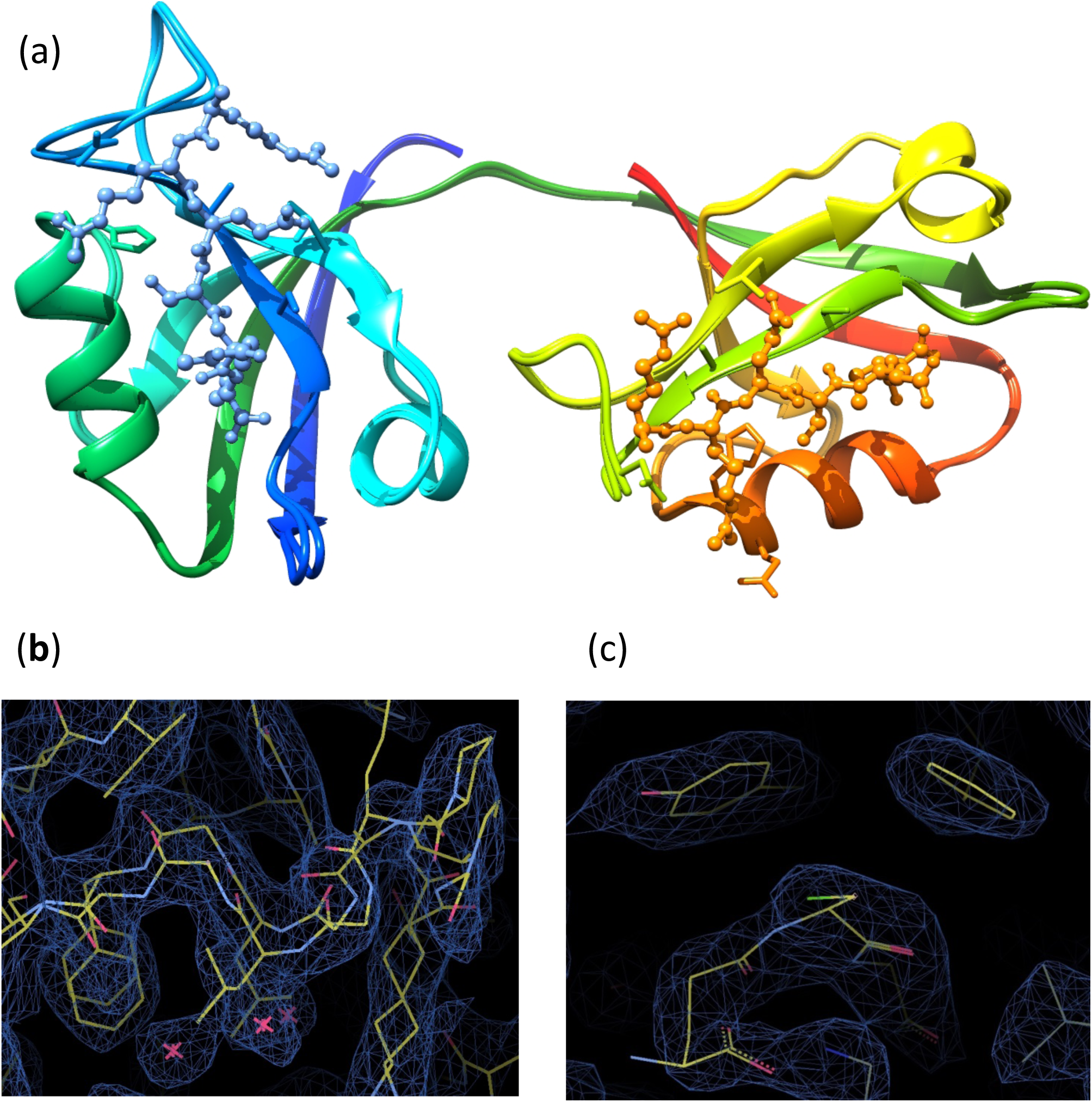
Crystal structure of PDZ1-2. The crystal structure of Apo: PDZ1-2 and RRESEI: PDZ1-2 are shown overlaid in (a), the PDZ1-2 double domain is represented in a similar way to Figure 1, with the associated RRESEI ligands shown in ball-and-stick form. The protein is coloured in a rainbow fashion from N-terminus to C-terminus with PDZ1 in blue-green and PDZ2 in yellow-red, generated with UCSF Chimera. Panel (b) shows the dual conformation of GLGF motif of PDZ2 in apo:PDZ1-2 and panel alongside chicken-wire 2Fo-Fc density (contour at 0.1 e^-^Å^-3^) (c) the 2Fo-Fc density assigned to GSH from apo:PDZ1-2 (0.2 e^-^Å^-3^). In b & c the model is shown in stick form with bonds to C, N, O, S in yellow, blue, red and green, generated with Coot (Emsley et al, 2010).

For RRESEI:PDZ1-2 the binding cleft of both PDZ domains is ordered and each binding site shows electron density for the RRESEI peptide. All of the ligand residues could be fitted, but the two Arg residues had markedly weaker electron density indicating that alternate conformations of these residues could be present. Similar interactions between the bound peptide and the clefts of both PDZ1 and PDZ2 are seen: The main chain amide linkages of the Ser residue of the incoming RRESEI peptide make H-bonds with the βB strand of both PDZ domains. A hydrogen bond is formed between the Ser alcohol and a His side chain (PDZ1:130, PDZ2:225) on the αB helix. Detailed examination of the electron density maps in the later stages of refinement of RRESEI:PDZ1-2 revealed weaker electron density for the C-terminal residue of each ligand peptide, with negative difference electron density enveloping the terminal carboxylate in difference electron density maps. This was interpreted as an alternative conformation of this part of the bound ligand, and was modelled with two alternative conformations of the C-terminal (-X_-1_-Φ_0_) -EI residues: One conformation has the Ile side chain buried in the binding cleft and the carboxyl terminus of the peptide associating with the GLGF motif, as seen in the structures of other Ligand bound Type I PDZ domains. The second conformation has the side chain of the Glu residue lying along the binding cleft with the Ile side chain making an interaction with Ile100/Ile195 in PDZ1/PDZ2 respectively. This model maintains a carboxylate function close to the GLGF motif of each PDZ domain for each alternate –EI conformation, whilst accounting for the observed features of the electron density. The model has a 0.5% lower R-free value (Brunger, 1992) compared to a single RRESEI conformation model (with each Ile sidechain buried in the cleft).

For Apo:PDZ1-2 the PDZ1 domain binding cleft is ordered but the PDZ2 domain shows disorder in both the βA-βB loop and αA helix regions. Weaker and more diffuse electron density is encountered for these segments which is consistent with disorder around the PDZ2 peptide binding cleft, including the GLGF motif. The αA helix links directly to the GLGF motif in PDZ2 within the binding cleft through main-chain hydrogen bonds between Leu 170 O and Ala 200 (O-N distance of 2.9Å) and Gly 171 and Ile 195 (O-N distance of 2.9Å). The electron density of the apo:PDZ2-βA-βB loop can be fitted with a dual conformation (Figure 2(b)). The two conformers are (*A*), a conformation similar to the RRESEI:PDZ1-2-βA-βB loop; and (*B*) a conformation similar to that found in the βA-βB loop of the syntrophin PDZ domain in the crystal structure of the neuronal Nitric Oxide Synthase/Syntrophin PDZ heterodimer (Hillier et al, 1999) (PDB ID:1qav). These two apo:PDZ2-βA-βB loop conformations were assigned the same occupancy but after refinement the residual B-factors were systematically lower for the RRESEI:PDZ1-2 -like conformation indicating a higher occupancy for this conformer.

In both Apo:PDZ1-2 and RRESEI:PDZ1-2 an additional electron density distribution was found adjacent to residues F119 and Y147. In both crystal structures the S-shaped electron density resembles a short peptide and has no connectivity linking it to the PDZ1-2 model: N and C-terminal residues (respectively GPNGT- and –SNA) from the tag-cleaved protein are not seen in the electron density maps. The location of the S-shaped electron density was distant from the C or N-terminus of any PDZ1-2 model in the crystal lattice. The electron density could not be accounted for satisfactorily by fitting the PEG crystallization precipitant for either Apo:PDZ1-2 or RRESEI:PDZ1-2. The cofactor reduced Glutathione (L-γ-glutamyl-L-cysteinyl-glycine, GSH) was used in all but the final protein purification step at milli-molar concentration (see Methods). A common association motif for GSH is the interaction of an amide plane from GSH with an aromatic residue side chain from the protein - for example PDB ID: 5bqg (Schiffler et al, 2016). The S-shaped electron density was fitted effectively with a single conformation of GSH with amide plane – aromatic stacking interaction with both F119 and Y147 (Figure 2(c)).

Differences between the individual PDZ1 and PDZ2 structures in apo:PDZ1-2 and RRESEI:PDZ1-2 are therefore limited to local changes in the vicinity of the peptide ligand binding site. The local disorder in apo:PDZ1-2 around the peptide biding site of PDZ2 and the αA helix is not seen in RRESEI:PDZ1-2 when the RRESEI peptide occupies the binding site. This shows that the peptide ligand binding site of PDZ2 is sensitive to the presence of the RRESEI ligand and that this is communicated to the αA helix. The peptide ligand binding site is linked to the αA helix via hydrogen bonds between the GLGF motif in the binding cleft and the helix in both PDZ1 and PDZ2 domains.

### Isothermal Titration Calorimetry

The crystal structure of RRESEI:PDZ1-2 shows that similar non-covalent interactions are observed for RRESEI association with both PDZ1 and PDZ2 binding clefts. A comparison of the crystal structures of apo:PDZ1-2 and RRESEI:PDZ1-2 indicates an ordering of the GLGF loop and αA helix regions of PDZ2 after RRESEI association. In contrast the PDZ1 binding cleft is the same in both structures. To further inform the observed differences in the association of ligand with PDZ1 and PDZ2, the thermodynamic changes on association of PDZ1, PDZ2 and PDZ1-2 domains with RRESEI were each measured using Isothermal Titration Calorimetry (ITC) (see Methods).

The thermogram of the PDZ1-2 and PDZ2 ITC experiment each showed exothermic peaks at every ligand injection. The thermodynamic components of binding extracted from these data indicated a negative enthalpic contribution (ΔH<0) to the free energy of binding, alongside a negative entropic contribution (ΔS<0). For the PDZ1 domain the thermogram was more complex showing exothermic binding initially, with a smaller endothermic binding change apparent close to saturation (Figure 3). The concentration of binding sites in the ITC experiment appeared considerably higher for PDZ1 and thermodynamic components for this domain indicate a negative enthalpic contribution (ΔH<0) and a positive entropic contribution (ΔS>0) (Figure 3.).

**Figure 3.**
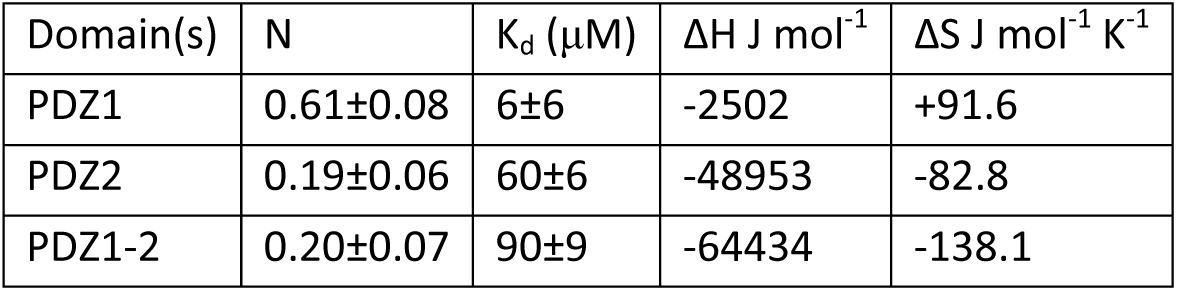
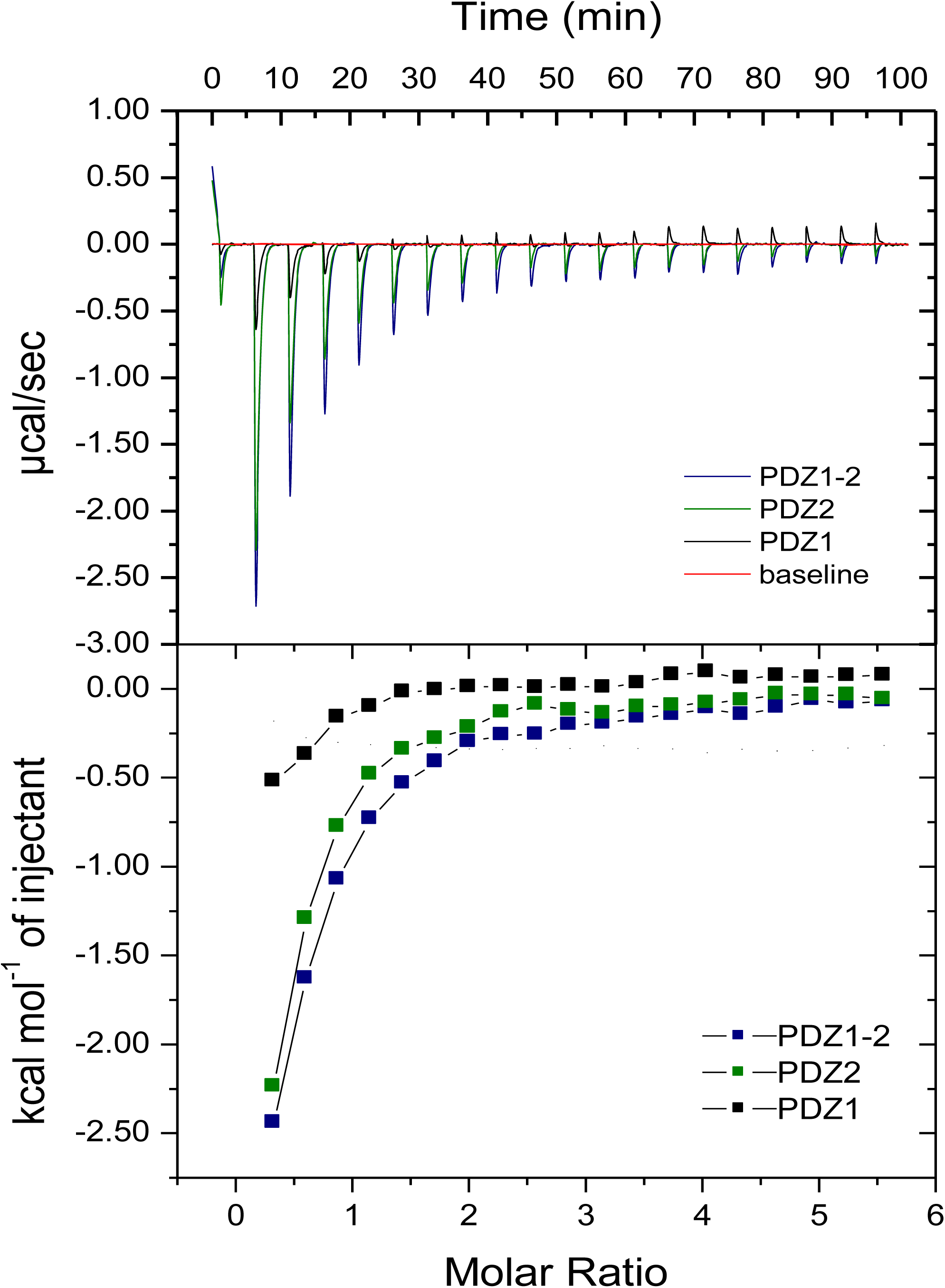
Isothermal Titration Calorimetry of the Association of PDZ1, PDZ2 and PDZ1-2 with the RRESEI ligand. ITC traces (top) and saturation analysis (bottom) for PDZ1, PDZ2 and PDZ1-2 each titrated with the RRESEI ligand are shown overlaid. The heat change on successive injections of ligand is shown in the top panel buffer heat of dilution subtraction (baseline in red). The corresponding binding isotherm with fitted curve is shown in the bottom panel. The value of the number of sites per molecule N, overall enthalpy change (ΔH) and equilibrium dissociation constant (K_d_) obtained from the top and bottom traces along with the entropy change (ΔS) derived are given in the accompanying table.

The equilibrium dissociation constant was measured at 6±6μM and 60±2μM for PDZ1 and PDZ2 respectively. The K_d_ values obtained for the affinity between the individual PDZ domains and the RRESEI peptide are within the range of those found for similar peptides binding to PDZ domains in the literature (Wiedemann et al, 2004). For the PDZ1-2 double domain two individual affinities could not be distinguished in the ITC measurement and an overall value of K_d_ of 90±9μM was obtained assuming a single class of sites. This value is high compared to the values of K_d_ obtained for the individual domains.

In the transition from Apo to RRESEI bound state a number of changes occur. The displacement of water from the binding cleft, formation of non-covalent PDZ domain-peptide interactions and the reduction in the degree of RRESEI structural freedom would appear to be similar for both PDZ1 and PDZ2 binding sites. The enthalpic component of the binding energy is mainly due to the formation of non-covalent interactions between RRESEI and the PDZ binding site. These interactions are similar for both PDZ1 and PDZ2 sites as seen in the RRESEI:PDZ1-2 crystal structure. However a large difference in the values of ΔH and ΔS is seen for isolated PDZ1 and PDZ2 domains binding to RRESEI.

For PDZ2 a net entropy decrease on RRESEI binding is observed. This is consistent with the ordering of the PDZ2 domain binding cleft on interaction with the peptide. This is seen when comparing the structures as the disordered βA-βB loop and αA helix regions of apo:PDZ1-2 crystal assume a single conformation in RRESEI:PDZ1-2.

The binding of RRESEI to PDZ1 arises from the concerted contribution of an entropy increase and an enthalpy decrease. In this case the interactions are formed between the RRESEI ligand and the ordered binding site seen in both the apo:PDZ1-2 and RRESEI:PDZ1-2 crystal structures. The additional endothermic contribution seen in the thermogram would tend to reduce the exothermic heat change at each ligand injection. Relative to RRESEI binding to PDZ2, the effects of the additional endothermic contribution seem to be a foreshortening of the saturation characteristic (leading to an increase in both number of binding sites N, and in affinity) along with a reduction in the integrated exothermic peak values (leading to a smaller change in enthalpy). This is reflected in the calculation of RRESEI:PDZ1 binding parameters: K_d_ and N from the saturation curve, and ΔH from the integration of heat changes. The entropy change ΔS is a derived parameter (obtained via K_d_ and ΔH), therefore a systematic change in all of thermodynamic parameters for RRESEI:PDZ1 association arises from the additional endothermic contribution observed in the case pf PDZ1. Therefore the difference in derived entropy changes and affinities between PDZ1 and PDZ2 can be accounted for. However the high value of the overall K_d_ for the RRESEI association with PDZ1-2 compared with the individual domains, and the source of additional endothermic element seen in the PDZ1 titration with RRESEI require explanation.

### Small angle X-ray scattering

The single conformation of PDZ1-2 in the crystal structures reported here accommodates binding of the RRESEI peptide ligand to both PDZ1 and PDZ2 domains. This conformation is different from those seen in the earlier PDZ1-2 crystal structures. The gross form of PDZ1-2 is therefore variable and may depend upon the interaction with the peptide ligand. To explore this variation the PDZ1-2 domain was examined in solution using Small angle X-ray Scattering (SAXS) in both the apo state and in the presence of a 10-fold molar excess of RRESEI. GSH was assigned in the crystal structures. GSH is present in the cell and GSH possesses both thiol and carboxylate functions in common with type I PDZ ligand sequences. GSH association with PDZ1-2 was also explored using SAXS. Data were collected on unfractionated samples and on samples fractionated using an in-line Size exclusion column prior to the measurement of scattering (see Methods). For each set of SAXS measurements, matching scattering data on apo:PDZ1-2 was collected. All data sets are summarized in Supplementary Table 2.

For the fractionated SAXS experiments an absorption chromatogram at a wavelength of 280nm was recorded immediately prior to exposure with X-rays. A scattering profile was then recorded for one or more fractions. In contrast to the preparative Size exclusion chromatography step (see Methods), a broader highly-absorbing peak was observed in these 280nm chromatograms preceded by a small peak/shoulder. The precise form of this main peak varied between sample injections, with flat top; unresolved doublet; or a single peak with a shoulder being observed. Where R_g_ analysis was carried out for multiple fractions (June ’16 data) the values of R_g_ obtained showed no significant variation across the main peak.

For the analysis of the SAXS emphasis was placed on the apo:PDZ1-2 and RRESEI:PDZ1-2 data sets collected on unfractionated samples at the DESY P12 beamline, these data sets extend to the highest resolution (q≤0.48Å^-1^) and have the lowest noise levels (See Supplementary Table S2; June ’15 data). Subsequently the GSH:PDZ1-2 and fractionated SEC-SAXS data sets were fitted using the formalism derived from the analysis of this higher resolution interval data.

The apo:PDZ1-2 and RRESEI:PDZ1-2 data showed evidence of both inter-particle association effects and underlying multiple conformations of PDZ1-2. The level of these two features differed according both the concentration of the PDZ1-2 protein and the presence of the RRESEI peptide ligand. For initial analysis, SAXS data projected to infinite dilution was used to ameliorate the presumed effects of inter-particle interference (see Methods). Dummy atom modelling of the SAXS data was limited to a truncated resolution range and gave a dumbbell like shape but with additional envelope features (Supplementary Figure S1.).

Ensemble model analysis based upon linked PDZ1 and PDZ2 domains was undertaken to account for variation in the gross form of PDZ1-2 (Bernado et al, 2007). The SAXS curve is very sensitive to the separations of protein domains in the sample and the ensemble analysis gave a clear indication that there were different domain separations represented in the SAXS data (see Methods). The presence of two or more particular conformations of PDZ1-2 was seen in contrast to a continuum of conformers like those derived from NMR.

An alternative explanation for the distributions seen in the ensemble analysis is the formation of inter-molecular complexes. A very compact structure (R_g_ ~21.2Å) consistently assigned in the ensemble analysis of Apo:PDZ1-2 is not represented by the crystal structure reported here or any of the preceding crystal structures (Bach et al, 2012; Sainlos et al, 2011). This domain configuration could arise from the close approach of two PDZ domains from separate PDZ1-2 molecules occurring on the formation of oligomers. Should these inter-molecular interactions be spatially specific in nature their effects would not be removed by projection of SAXS curves to infinite dilution.

### Oligomer Model Construction

Crystal contacts and PDZ1-2 conformations drawn from existing crystal structures were used to generate oligomers for fitting the SAXS data. The rationale for this approach was that at the ultra-high protein concentrations represented in protein crystals, inter-molecular interactions would appear as crystal contacts. Dimeric and trimeric oligomers of PDZ1-2 were constructed using: (i) The extended 3zrt:PDZ1-2 monomer, with (ii) two crystal contacts present in the 3gsl:PDZ1-2 crystal lattice, between αB(PDZ2):βD-βE(PDZ1) and αA(PDZ2): βB-βC(PDZ1) (shown in Supplementary Figure S2). These components give rise to dimer and trimer forming interactions between 3zrt:PDZ1-2 monomers and using these symmetry components more extended arrays of PDZ1-2 can be constructed (see Methods).

There are a finite number of ways that single molecules can arrange themselves in extended repeating arrays in 3-dimensions (Hahn, 1983). Therefore the assignment of a Spacegroup to the oligomeric structures encountered for PDZ1-2 was investigated. A cubic I2_1_3 Spacegroup with a unit cell parameter |**<u>a</u>**|= 148Å could be assigned to the oligomers of PDZ1-2 in an unambiguous manner (see Methods). The αB(PDZ2):βD-βE(PDZ1) and αA(PDZ2): βB-βC(PDZ1) interfaces are properties of this arrangement and a further PDZ1:PDZ1 interaction is predicted by the molecular packing. Various oligomers can be reproduced by selecting the appropriate combinations of symmetry operations from this I2_1_3 “scaffolding Spacegroup”.

### Order of Oligomer Assembly

The formation of any protein-protein complex is governed by the abundance of the assembling components and the affinity of the components for one another. In the scaffolding Spacegroup each PDZ1-2 monomer makes two interactions of the form αB(PDZ2):βD-βE(PDZ1), two of αA(PDZ2):βB-βC(PDZ1) and one PDZ1:PDZ1 interaction. There is no reliable independent measurement of affinity for these three individual interactions. The interactions are all non-covalent with hydrogen bonding and ion pair interactions dominating each interface. Therefore in the analysis of assembly it was assumed that each of these interactions affinities are similar in magnitude, whilst considering the formation of oligomers as the overall concentration of PDZ1-2 increases. The order of assembly is then determined via the principle of avidity: A meta-stable complex occurs when multiple (avid) interactions form between binding partners because of the factorial increase in affinity. In other words interactions at two or more separate sites would need to be broken at the same time for a complex so formed to dissociate.

A PDZ1-2 dimer may form through any of the interactions listed above, with few restrictions on the gross conformation of PDZ1-2. If a 3zrt:PDZ1-2 like conformation is adopted by the binding partners two αB(PDZ2):βD-βE(PDZ1) interactions can form at the same time (Figure 4. (a)2e). This “double” dimer would therefore be meta-stable over the other pairwise interactions due to avidity. This dimer/monomer mixture can be refined against SAXS data (see Methods), with good agreement achieved for lower concentrations of PDZ1-2 (SEC-SAXS or data extrapolated to zero concentration).

**Figure 4.**
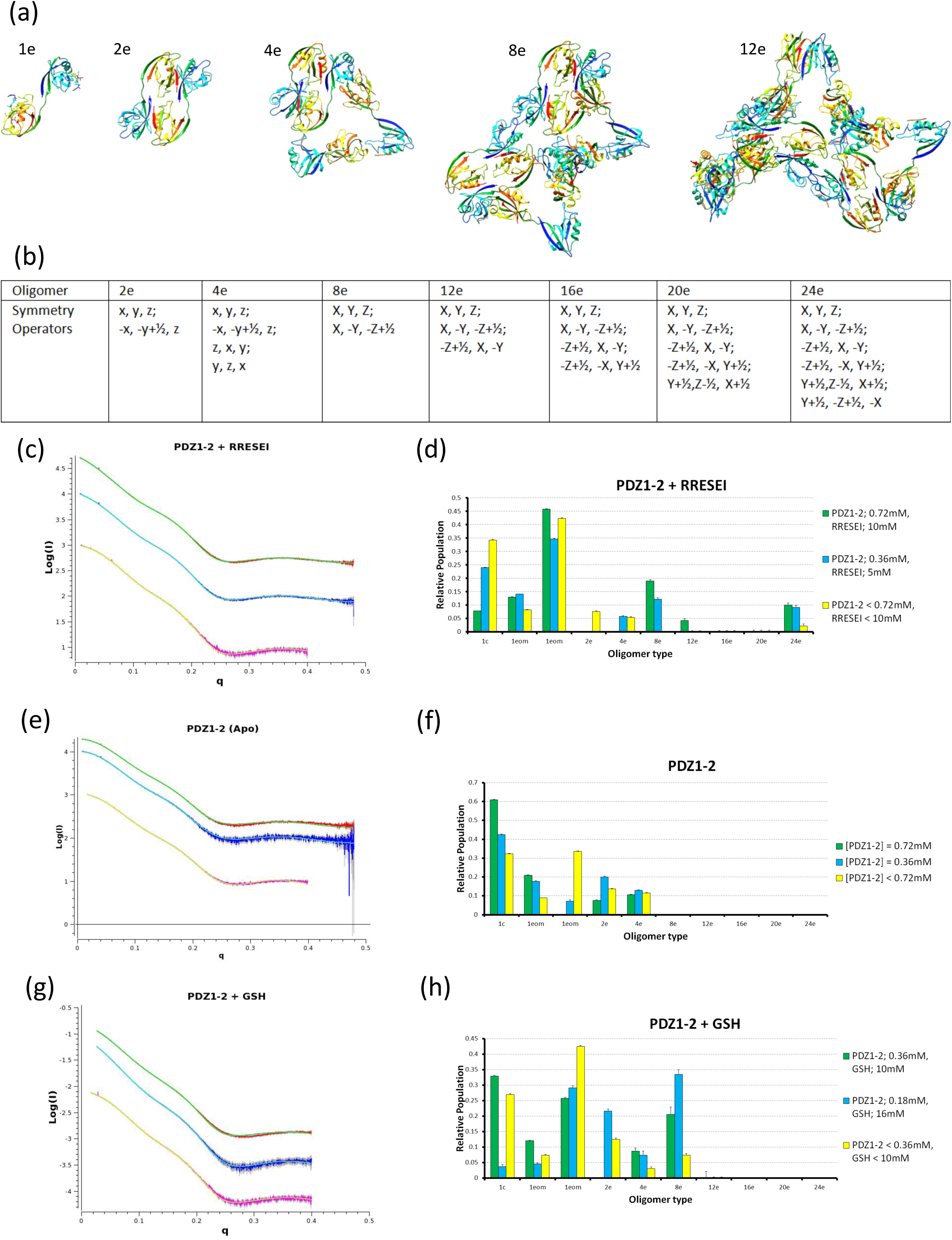
Oligomer fitting of SAXS data. Selected PDZ1-2 oligomers 1e, 2e, 4e, 8e and12e are shown in (a) where “e” denotes the extended 3zrt:PDZ1-2 conformation. The symmetry operations drawn from the scaffolding Spacegroup required for the construction of the PDZ1-2 oligomers used in fitting are given in (b). Transformations applied to the fractional coordinates of the PDZ1-2 monomer ((a).1e) given in lower case (x, y, z), and those applied to the meta-stable tetramer ((a).4e) in capital letters. Fitted SAXS curves are shown alongside histograms of the oligomer populations in (c)-(h). Scattering curves are plotted as Log(I) versus q throughout (where I is the scattering intensity and q is momentum transfer). High concentration data curves are shown in red with fitted curves in green, dilutions in blue with fitted curve in cyan and SEC-fractionated samples in magenta with fitted curves in yellow. A multiplication factor has been applied to the raw data in some cases to separate curves along the ordinate (Log(I)) axis. The Histograms show the relative populations of various oligomers determined in the fitting analysis. The populations were determined by multiplying the volume fraction assigned by the OLIGOMER program (Franke et al, 2017; Konarev et al, 2003) by the oligomer number and then renormalizing all fractions to sum to 1. Hence the columns show the proportion of PDZ1-2 molecules assigned to each fraction. The “1c.” oligomer shown on the abscissa maps to compact PDZ1-2, the 2 “eom” oligomers to uncoupled versions of PDZ1-2 derived from analysis of RRESEI:PDZ1-2 with the EOM program (Bernado et al, 2007), 2e-24e are drawn from the scaffolding Spacegroup.

Proceeding from this meta-stable dimer a further complex could be formed by either an αA(PDZ2):βB-βC(PDZ1) or a PDZ1:PDZ1 interaction. Either a dimer:monomer interaction or a dimer:dimer interaction would be possible and the affinity of these would be the same. Therefore a dimer:monomer complex would be favoured initially until the concentration of dimers exceeded the concentration of monomers. There would be few restrictions on the conformation of the associating monomeric PDZ1-2.

The addition of another monomer to this dimer:monomer complex to form a tetramer could again occur through any of the aforementioned interactions. In one particular case, where the preceding dimer:monomer interface was formed via αA(PDZ2):βB-βC(PDZ1) interaction, a further monomer may add forming two αA(PDZ2):βB-βC(PDZ1) interactions; again subject to the adoption of a 3zrt:PDZ1-2 like conformation for all PDZ1-2 copies. This tetramer complex (Figure 4(a)4e) would be meta-stable since in the oligomer so formed each PDZ1-2 is involved in at least two interactions. The meta-stable tetramer complex has three copies of PDZ1-2 related by a 3-fold rotation axis and two copies related by a 2-fold rotation axis. This configuration can be refined against SAXS data as mixture with the PDZ1-2 monomer, good agreement is achieved for lower concentrations where the RRESEI ligand is present. The angle between these 3-fold and 2-fold symmetry elements is variable in the resulting refined models with a larger angle (≈ 90°) seen for refinement with data collected on more dilute samples. A consequence of a large angle is that the addition of more monomers to the tetramer via αB(PDZ2):βD-βE(PDZ1) interactions would be sterically hindered.

In the scaffolding Spacegroup the angle between the 3-fold and 2-fold symmetry elements is fixed at 54.7°. Thus a particular conformation of the meta-stable tetramer is required for assembly into even higher order structures, in a similar way to the requirement of the monomeric form of PDZ1-2 to adopt the 3zrt:PDZ1-2 conformation for complex formation. When the scaffolding Spacegroup tetramer configuration is used two tetramers can associate to form an octamer by forming two αB(PDZ2):βD-βE(PDZ1) interactions. In one of the three possible configurations of this octamer an additional PDZ1:PDZ1 contact is also made. This particular configuration would therefore be favoured as three interactions are formed on assembly (Figure 4(a)8e). Subsequent additions of tetramers each making three interactions may then be made for 12 (Figure 4(a)12e), 16, 20 and 24-mer oligomers, the latter corresponding to the components of the unit cell of the scaffolding Spacegroup.

### SAXS data fitting

Oligomers drawn from the scaffolding Spacegroup were assessed for fitting the SAXS curves (Figure 4. (c)-(h)). As variation in the structures of monomeric PDZ1-2 was also evident this was captured by the inclusion of three representative monomer structures – the compact version of PDZ1-2 reported here, plus two PDZ1-2 structures extracted from a representative ensemble analysis run on SAXS data extrapolated to zero concentration using the EOM program (Bernado et al, 2007) (see Methods). The EOM derived monomers were similar in form to structures found in 3gsl:PDZ1-2, or 2ka9:PDZ1-2, having no direct, non-covalent, intra-molecular PDZ1-PDZ2 contacts. Agreement with the SAXS profile was improved, in particular for RRESEI:PDZ1-2, at all concentrations by the inclusion of oligomers derived from the scaffolding Spacegroup. Volume fractions assigned to the various oligomers for apo:PDZ1-2 and RRESEI:PDZ1-2 are shown in Figure 4(d,f). The scattering data collected in the presence of GSH was fitted simply through direct application of this model. In the case of GSH:PDZ1-2 the relative concentration of GSH and PDZ1-2 vary for each Scattering experiment (see Methods): Here the higher order oligomers are associated with the highest GSH:PDZ1-2 ratio (Figure 4(h)). The methodology deployed to interpret the scattering data is compelling because relatively few degrees of freedom are required to describe the complete structural model, and variation of oligomer populations with PDZ1-2 concentration is readily accounted for. Compared to the dummy atom modelling with data projected to infinite dilution the values of Χ^2^ are similar: For RRESEI:PDZ1-2 Χ^2^ values of 8.74, 2.71 and 3.71 were obtained for concentrated, diluted and SEC data respectively, the equivalent values for Apo:PDZ1-2 are 6.31,1.62 and 2.67 and for GSH:PDZ1-2, 1.41, 0.33 and 4.21. The Oligomer based fitting is an improvement as scattering data over the complete momentum transfer interval recorded is included (Figure 4(c, e, g)).

For all of the SAXS experiments reported here, lower concentrations of PDZ1-2 arise from a higher concentration sample, either through dilution, or for the fractionated data via diffusion during passage down a Size exclusion column (see Methods). We assume that the timescales of the biophysical experiments here are long enough such that equilibrium conditions pertain. Some hysteresis in the formation of assemblies appears to be present as the larger oligomers persist after dilution. This may reflect a difference in assembly through the progressive formation of larger and larger complexes over dis-assembly via the dissociation of smaller components such as monomers. Data fitting with OLIGOMER was restricted to the metastable complexes identified in the considerations of avidity (monomer, dimer, tetramer, octamer, 12, 16, 20 and 24-mer oligomers). In the case of apo:PDZ1-2 monomer dimer and tetramer oligomers are sufficient to account for the scattering (Figure 4(f)). In the case of RRESEI:PDZ1-2 oligomers extending up to a 24-mer are needed to account for the observed scattering profile (Figure 4(d)). Fitting of SAXS curves was limited to oligomers of up to 24 copies of PDZ1-2 as this corresponds to the number of general equivalent positions in the unit cell of the I2_1_3 Spacegroup (Hahn, 1983). Some marginal improvements in the agreement of the curve with the data were observed if larger oligomers were included. However the inclusion of more than one unit cell of a three dimensional lattice model would imply some constructive interference in the Scattering: As such this may break assumptions inherent in the methodology used to generate calculated SAXS curves which are fitted. The GSH:PDZ1-2 data requires fewer oligomers drawn from the scaffolding Spacegroup with good agreement from oligomers of up to 8 copies of GSH:PDZ1-2 (Figure 4(h)).

### Compatibility of RRESEI and GSH ligands with the oligomers

The thermodynamic parameters extracted from the ITC analysis of PDZ1-2:RRESEI binding are not simply related to their counterparts extracted for isolated domains. This suggests that either binding-site interference or intermolecular interactions may be at play. The crystal structures of PDZ1-2 do not support any mechanism of direct site interference for a short peptide like RRESEI, whereas the SAXS measurements show that the RRESEI enhances the formation of PDZ1-2 oligomers. Despite the presence of the PDZ1 domain in PDZ1-2 the endothermic peaks observed near-saturation for PDZ1 are not seen in titrations of PDZ1-2 with RRESEI. Association of RRESEI with PDZ1-2 is also correlated with a lower population of the compact form of PDZ1-2. The transition from compact to more extended PDZ1-2 increases the number of conformations that can be adopted by PDZ1-2; alongside the replacement of an intra-molecular interaction with a solvent interaction. The increased number of gross conformational states would lead to an increase in entropy +ΔS and the exchange of interactions a small change in enthalpy ΔH. According to the SAXS analysis presented here the αB(PDZ2):βD-βE(PDZ1) double dimer forms in the apo case but the population is enhanced by the addition of RRESEI. This configuration would be favoured over a direct interaction between RRESEI:PDZ1 domains. The formation of the double-dimer fixes the conformation of PDZ1-2 alongside the formation of non-covalent interactions at the two PDZ2:PDZ1 interfaces (giving thermodynamic contributions -ΔS and -ΔH). The additional factors leading to RRESEI:PDZ1-2 oligomerization therefore make diverse thermodynamic contributions. These additional interactions could readily account for the differences between the separate PDZ domains and the double domain seen in the ITC titrations with RRESEI.

The RRESEI ligand can be accommodated within the αB(PDZ2):βD-βE(PDZ1) interface after the adjustment of side chain rotamers of aromatic residues Y63 and Y147 of PDZ1-2. The ligand peptide may then participate in a continuous β-sheet with RRESEI intercalated between the βA strand of PDZ1 (forming parallel β-sheet interactions) and the βB strand of PDZ2 (forming anti-parallel β-sheet interactions). The GSH molecule identified in the Apo:PDZ1-2 and RRESEI:PDZ1-2 structures associates with Y147 and F119. A GSH molecule at this site on PDZ1 would be presented to the PDZ2 binding cleft on formation of the αB(PDZ2):βD-βE(PDZ1) interface. The GSH contains thiol and carboxylate GSH can readily form interactions within the binding cleft. The GSH site found the PDZ1-2 crystal structures is therefore likely to contribute to the affinity of the αB(PDZ2):βD-βE(PDZ1) interface, but GSH would be excluded from the interface if a peptide ligand was present in the PDZ2 cleft. Therefore both GSH and RRESEI are able to knit the αB(PDZ2):βD-βE(PDZ1) interface together.

The other contact used in defining the I2_1_3 scaffolding lattice is αA(PDZ2): βB-βC(PDZ1). There is no direct involvement of binding clefts in this interface. The relationship between the stability of the αA region in PDZ2 and the association of RRESEI at the PDZ2 cleft indicates that ligand recognition may be an indirect factor in the formation of this contact.

A direct interface between copies of PDZ1 is predicted within the I2_1_3 scaffolding lattice. An interface of this type can form between isolated PDZ1 domains. The SAXS analysis indicated that the formation of this interface is important for larger oligomers like those required for fitting RRESEI:PDZ1-2: In turn suggesting that the binding of RRESEI enhances the formation of the interface. This would explain the unusual behaviour of the PDZ1 domain in ITC where an additional endothermic heat change is resolved close to saturation with the RRESEI ligand: The additional heat change observed being due to the association of RRESEI:PDZ1 domains with one another. In the ITC experiment the PDZ1 domain is successively diluted in the measurement cell by the addition of aliquots of the RRESEI solution. Thus any association of PDZ1 domains must be RRESEI mediated.

Interfaces between copies of peptide bound PDZ domains have been seen in crystal structures, for example PDB ID 1oby (Kang et al, 2003). The alternative orientation of RRESEI with the C-terminus emergent from the PDZ1 ligand binding cleft seen in refinement of the RRESEI:PDZ1-2 crystal structure appears to be compatible with this interface in the scaffolding Spacegroup. When this orientation is present additional interactions are made at the interface including the insertion of Asn72 between the bound ligand and αB helix of PDZ1. However as the scaffolding arrangement is effectively determined by SAXS analysis, which is limited to a lower resolution, structural details of this type should be treated with caution.

PDZ1-2 oligomers in the presence of GSH alone are more limited in extent than those in the presence of RRESEI. The influence of GSH seems to be primarily in the αB(PDZ2):βD-βE(PDZ1) interface. Using the scaffolding Spacegroup it is possible to devise octamer and dodecamer assemblies of PDZ1-2 tetramers which do not have a PDZ1:PDZ1 interaction (a dodecamer can be formed from tetramer symmetry operations in similar manner to figure 4(b) X, Y, Z; Y,-Z,-X+1/2; -X+1/2, Y, -Z). The shorter GSH peptide ligand may not significantly enhance the PDZ1:PDZ1 interface predicted in the scaffolding Spacegroup and this may explain why oligomers are more limited in extent in the presence of GSH.

## Discussion

The preparation of sub-cellular neuronal compartments such as PSDs for imaging or extraction of proteins often necessitates harsh treatments such as mechanical disruption, multiple centrifugation steps and extensive dilution (Carlin et al, 1980). A recent report also indicates that detergent-specific effects can occur when isolating component oligomers from PSD fractions (Lautz et al, 2019). So many of the non-covalent interactions revealed here are likely to be compromised in sub-neuronal isolates. Therefore the approach of assembling component elements taken in this study has the power to reveal new information. In their dissection of the clustering properties of PSD-95 Hsueh *et al.,* noted that “A rafting mechanism for clustering of multimeric channels … could still be accomplished by a single PDZ domain per PSD-95 monomer if PSD-95 self-associates to form multimers”. The work here shows that extensive association of PSD-95 can occur but this is acutely sensitive to the binding ligand. Also since the association is driven by relatively weak non-covalent interactions a high concentration of PSD-95 is required.

### The role of the peptide ligand

A significant feature of the Apo:PDZ1-2 crystal structures reported here is the disorder of the PDZ2 cleft and the PDZ2 αA helix. In contrast for 3gls:PDZ1-2 the vacant PDZ2 ligand binding site is ordered for both copies of PDZ1-2 (this is also true of 3zrt:PDZ1-2 although lower resolution is a factor for this structure). In both 3gls:PDZ1-2 and 3zrt:PDZ1-2 the binding cleft of each PDZ2 domain is involved in the αB(PDZ2):βD-βE(PDZ1) interaction with a PDZ1 domain from another PDZ1-2. This αB(PDZ2):βD-βE(PDZ1) inter-molecular contact may therefore stabilize the conformation of the vacant PDZ2 ligand binding site. In 3gls:PDZ1-2 inter-molecular contacts between Tyr 147 and His 225, and Glu 65 and Ser 173 are formed. In 3gls:PDZ1-2 the αA helix of PDZ2 is additionally stabilized by a crystal contact αA(PDZ2): βB-βC(PDZ1) with a second PDZ1-2 molecule.

The two bound RRESEI peptides in the RRESEI:PDZ1-2 structure show similar features including weaker electron density at their C-terminus, which is interpreted here with a dual conformation of the terminal EI residues of both of the bound peptide ligands. Both PDZ1 and PDZ2 peptide binding clefts appear ordered in RRESEI:PDZ1-2. The disorder of the C-terminal residues of the RRESEI ligand in the PDZ1 cleft seen in RRESEI:PDZ1-2 would appear to be an unexpected result given the ordered nature of the unoccupied PDZ1 site in apo:PDZ1-2. The alternate bound conformation of RRESEI lacks the hydrophobic group inserted into the cleft but forms a contact with a residue in a loop immediately preceding the αA helix.

The stability of the PDZ2 peptide ligand cleft is enhanced when inter-molecular contacts are made between the cleft, the adjacent αA helix or when a peptide ligand is bound. This indicates that there is likely to be a synergy between the conformation of the PDZ2 cleft/PDZ2 αA helix, and the presence of the peptide ligand. This can be mediated by the GLGF sequence which links these two sub-structures via non-covalent interactions.

The gross structures of 3gsl:PDZ1-2 is less compact than RRESEI:PDZ1-2 because the intra-molecular αA(PDZ1): βB-βC(PDZ2) interaction in RRESEI:PDZ1-2 is not formed in 3gsl:PDZ1-2. In the 3gls crystal structure the –ETMA ligand and the PDZ1 cleft are ordered and the Met residue forms a hydrophobic contact with Ile 100. There is no intra-molecular contact between the βB-βC loop of PDZ2 and the αA helix of PDZ1.This raises the possibility that a synergy between the binding cleft and the αA helix of PDZ1 also exists: The ligand peptide may induce a conformational change in the peptide binding cleft which is relayed to the αA helix by the GLGF region and a hydrophobic contact between the ligand and Ile-100 which in turn de-couples the αA(PDZ1): βB-βC(PDZ2) intra-molecular interaction.

Taken together these facets indicate that a subtle induced fit mechanism may be at work for the interaction of RRESEI with PDZ1-2. RRESEI interaction at PDZ1 induces conformational changes which in turn are relayed to the αA helix of PDZ1. An uncoupled conformation of PDZ1-2 results from the dissociation of the intra-molecular interaction. In contrast RRESEI peptide interaction at PDZ2 induces a single ordered conformation of the GLGF bearing cleft, which in turn is relayed to the αA helix of PDZ2 favouring the ordered conformation of this helix seen in RRESEI:PDZ1-2.

Some variation in the intra-domain separation of PDZ1 and PDZ2 has also been reported in NMR studies (Wang et al, 2009), and domain separation was enhanced in the presence of a ligand. Modelling of domain separation variation of this nature using ensemble model-fitting software (Bernado et al, 2007) was effective for SAXS data extrapolated to zero concentration where contributions from oligomers are smaller.

### The role of the conformation of PDZ1-2

The gross form of the PDZ1-2 domain is like a “dumbbell” and two peaks consistent with this structure are clearly evident in the Pair Distance distribution histograms derived from the scattering curves (see Supplementary materials). Underlying this overall form were systematic differences between both the scattering curves and their corresponding transformed pair distance distribution functions according to the concentration of PDZ1-2 and the presence of peptide ligands. Pair distributions of PDZ1-2 in complex with a peptide based inhibitor of PSD-95 clustering also showed a similar form (Bach et al, 2012): The pair distribution function of PDZ1-2 in presence of a monomeric inhibitor shown for this work appears to show a sharp peaks in P(r), which is qualitatively similar to that encountered in the RRESEI:PDZ1-2 case here (see Supplementary materials).

The population of the compact form of PDZ1-2 is low for the high concentration of RRESEI:PDZ1-2 as shown by the SAXS analysis (Figure 4(d)), with a small population of the compact form seen only for diluted RRESEI:PDZ1-2 (Figure 4(d)). Thus the RRESEI:PDZ1-2 crystal structure described here represents a minority conformer in solution, which may explain the requirement for seeding in the formation of RRESEI:PDZ1-2 crystals (the seeding promoting nucleation at lower RRESEI:PDZ1-2 concentration where there is a significant level of the compact monomeric form). In contrast in the apo case the compact form of PDZ1-2 is present at an increased concentration and the PDZ1-2 inter-molecular interactions are more limited (Figure 4(f)). This indicates a preference for the decoupled/extended conformation in the presence of bound ligand. A synergy between the association of RRESEI peptide at the PDZ1 binding cleft, the conformation of the αA helix of PDZ1 and the gross conformation of PDZ1-2 is a likely reason for this.

A major determinant in the formation of the I2_1_3 scaffolding lattice revealed here is the gross conformation of the PDZ1-2 domain. The trimer-containing oligomers which in turn lead to the cubic packing arrangement require the PDZ1-2 domain to adopt an extended (3zrt:PDZ1-2 -like) form. A first step in the establishment of this extended conformation is the decoupling of the compact form of PDZ1-2. The ligand PDZ2 interaction is therefore key to the trimer forming interaction (αA(PDZ2): βB-βC(PDZ1)) and in turn fundamental to the I2_1_3 scaffolding lattice identified here. Additionally a ligand sequence incorporated at the αB(PDZ2):βD-βE(PDZ1) interface may change the affinity of this interaction. The properties of residues “X” in the sequence –X_-3_–(Ser/Thr/Cys)_-2_-X_-1_-Φ_0_ from the partner channel may modulate the affinity of MAGUK scaffolding interactions. In the work described here the presence of a Glu residue at X_-1_ appears to be important in allowing an alternate peptide ligand conformation. Additionally the aliphatic side chain at Φ_0_ may have an influence on either the perturbation of the PDZ ligand binding cleft in the buried conformation, or the formation of a contact with the loop adjacent to the αA helix. High concentrations of MAGUK proteins are present in functional compartments of neurones alongside their partner receptors and channels. The NMDA receptor present in the PSD is assembled from combinations of four ε and ζ subunits (Hess et al, 1998), the ε subunits have a C-terminal sequence of –E_-3_-S_-2_-(D/E)_-1_-V_0_. Hence the NMDA receptor also possesses carboxylic acid bearing residues at the X_-1_ position and will have similar effects in PDZ1-2 as RRESEI.

### The PDZ1-2 domain forms the core of a MAGUK scaffold

The PSD-95 protein contains five modules only two of which are considered in the work reported here; hence the arrays shown may be affected by the other domains of the protein. The PDZ1-2 module of PSD-95 is bracketed by two long unstructured peptide linkers (Figure 1.1) which will allow relative freedom for the PDZ1-2 module to form interactions with other copies of PDZ1-2 close by. However the closest interaction partner for an isolated PDZ1-2 unit will be PDZ3. It is relevant therefore to examine whether the PDZ3 domain could form similar domain-domain interactions to those found in the PDZ1-2 scaffolding Spacegroup. The precise affinity of any PDZ:PDZ interaction will be governed by the amino acids forming interfaces including any associated ligand peptides, but compatible secondary structure elements also need to be in place. PDZ3 has additional secondary structure elements after the βF strand and also has a truncated βB-βC loop (Doyle et al, 1996). This means that PDZ3 could substitute for PDZ2 in the scaffolding Spacegroup (αB(PDZ3):βD-βE(PDZ1); αB(PDZ2):βD-βE(PDZ3) and αA(PDZ3):βB-βC(PDZ1) could all form). PDZ3 can form the (RRESEI:PDZ1):(RRESEI:PDZ3) interaction, however the domain could not form an αA(PDZ2):βD-βE(PDZ3) interaction. Therefore PDZ3 cannot fully substitute for PDZ1 in the scaffolding Spacegroup. Based upon this simple analysis PDZ3 is likely to integrate with the scaffold. As such the PDZ1-2 domain may form a core scaffold which can be decorated by PDZ3. It is important to note that in the scaffolding Spacegroup all copies of PDZ1-2 form two αB(PDZ2):βD-βE(PDZ1) interactions. The avidity due to this dual interaction means that the dual domain inter-molecular interaction is likely to displace any PDZ1-2:PDZ3 intra-molecular interaction on the close approach of two PDZ1-2 units.

The PSD-95 N-terminal linker has been found to be essential for clustering (Hsueh et al, 1997). The N-terminal linker requirement is likely due to membrane localization which is a factor in maintaining a high local concentration of PSD-95 near the membrane. There are variations in the N-terminal linkers with the second isoform of PSD-95 having a longer (98 residue) linker. In addition the closely related PSD93 MAGUK has been shown to be palmitoylated at the N-terminus (in Rat) (El-Husseini et al, 2000b) and may participate in the formation complexes similar to those found here.

The SAXS experiments indicate that the formation of extended oligomers of PDZ1-2 requires saturating concentrations of type I PDZ peptide ligands. These sequences are found in the cytoplasm at the ends of long unstructured regions at the C-termini of trans-membrane ion channels, this condition is also satisfied close to the surface of a membrane bearing a high density of such channels. Likely arrangements of PDZ1-2 oligomers are shown in Figure 5(a,b). Extended oligomers were observed in earlier work with the RRESEI containing the Kir2.1 cytoplasmic domain and full-length PSD-95 (Fomina et al, 2011) with both components in the solution phase. A PDZ1-2 oligomer coincident with the 3_1_ screw axis of the scaffolding Spacegroup could form the core of these structures as illustrated in Figure 5(c). In order to form a 2 dimensional net PDZ1:PDZ1 interactions are necessary to link within and between these extended oligomers (Figure 5(a,b)). As indicated in Figure 5(b) the PDZ1:PDZ1 interactions lie toward one side of the net shown. The ligands required to enhance the interfaces between those copies of PDZ1-2 are available close to the membrane surface.

**Figure 5.**
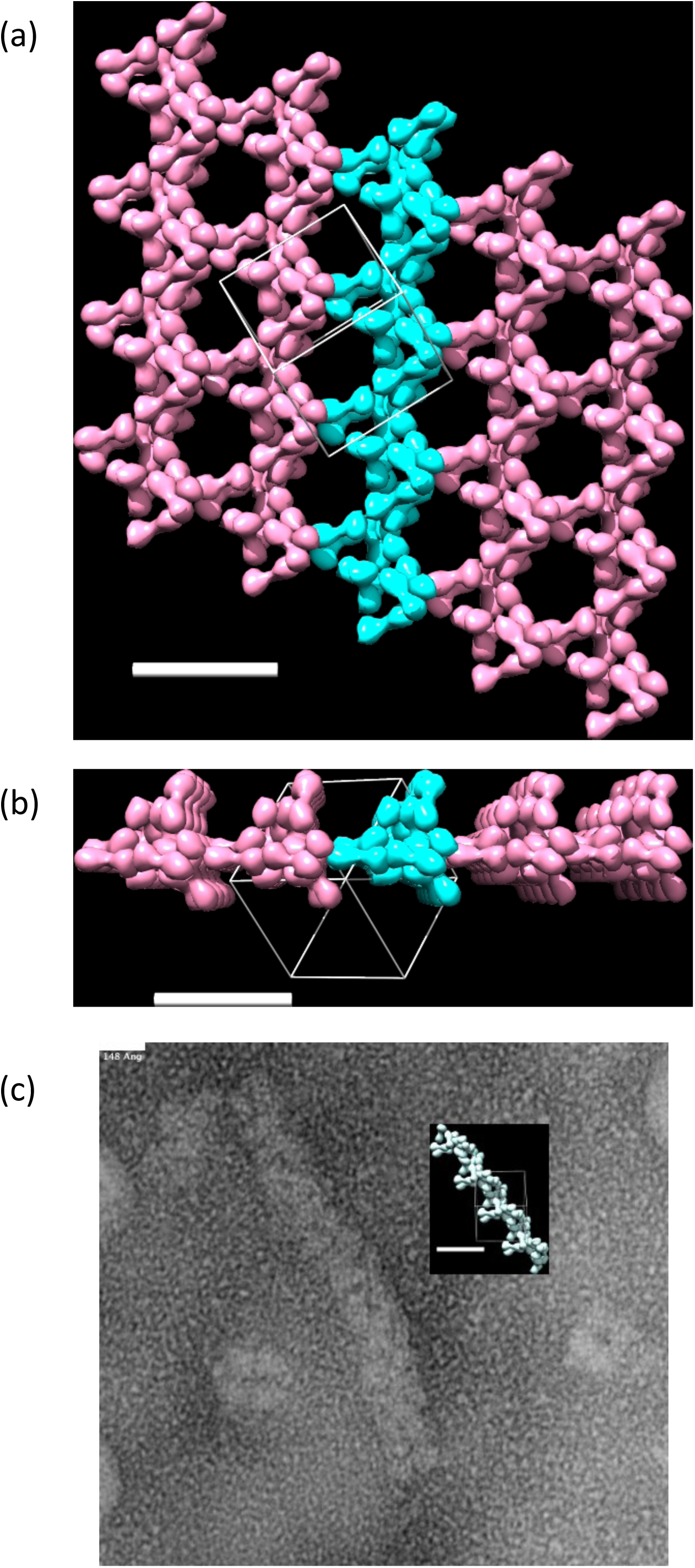
Higher order PDZ1-2 structures. Plan and end elevation views of a PDZ1-2 net are shown in (a) and (b) respectively, each PDZ1-2 domain is represented by a surface envelope. An oligomer coincident with the 3_1_ axis of the scaffolding Spacegroup is picked out in (a) and (b) and shown as a scaled inset in (c). All scale bars are 148Å and a unit cell outline for the scaffolding Spacegroup is included with the molecular representations. The electron micrograph shown in (c) was selected from the data for PSD-95 and Kir2.1 complexes (Fomina et al, 2011) and rendered with ImageJ software (Schneider et al, 2012), molecular representations are from UCSF Chimera.

### The scaffold Spacegroup has optimal properties

Scaffolds formed *in vivo* would therefore be subject to the high concentration of both ligand sequence and MAGUK, which is limited to a volume close to the membrane. This will allow the formation of a net of ligated PDZ1-2 but a deeper 3-dimensional array of PDZ1-2 cannot form. GSH is an indigenous cellular peptide, the SAXS work here shows that GSH has similar effects on PDZ1-2:PDZ1-2 association as the RRESEI ligand. These effects are evident at mM concentrations of GSH. GSH pervades the cytoplasm of cells at 0.5-10mM concentration (Wang & Ballatori, 1998). However GSH:PDZ1-2 oligomers are more limited in extent. Thus nets formed by means of ligated PDZ1-2 scaffolds may be maintained through the integration of GSH, but cannot be significantly extended by incorporation of GSH.

In common with all cubic Space groups I2_1_3 is isotropic (the same in three orthogonal directions) on the mesoscopic scale. Hence the packing arrangement dictated by the scaffolding Spacegroup could be preserved whilst following the membrane curvature seen in specialized structures such as synaptic boutons or cylindrical axons. Long bi-directional nets can readily be formed (Figure 5.) and these arrangements are compatible with the restrictions of peptide ligand availability close to a surface. The assembly based upon the meta-stable tetramers the assembly naturally follows the network of crossed 3_1_ screw axes which are parallel to “isotropic” body diagonal directions of the I2_1_3 scaffolding lattice (Figure 5 (a)). A net formed in this way has regularly spaced voids (Figure 5(a)) allowing space for other components to integrate with the scaffold. The receptors and channels organized by this scaffold would follow this executive organization with an underlying repeating unit of the order of 14.8nm determined by the unit cell of the I2_1_3 scaffolding lattice.

### Scaffold modulation

A compelling feature of the symmetry of the scaffolding arises if one considers breaking the precise symmetry of the PDZ1-2 double dimer (seen in Figure 4(a)2e). This asymmetry can be accommodated with a revised P2_1_3 scaffolding Spacegroup with an identical unit cell length. Thus as long as the residues involved in the αA(PDZ2):βD-βE(PDZ3) and PDZ1:PDZ1 contacts are compatible, the packing of heterogenous PDZ1-2 domains can occur in a similar way. Other MAGUK proteins are found alongside PSD-95 in synaptic fractions including SAP97 (Li et al, 2004). The PDZ1-2 regions of the canonical isoforms of PSD-95, SAP97, PSD93 and SAP102 have an amino acid identity of 72%. The PDZ1-2 region of SAP97 has an identity of 89% with PSD-95 but is not localized to the membrane via post-translational modification like PSD-95 (and PSD93). It has been shown that PSD-95 and PSD-93 can hetero-multimerize (Kim et al, 1996) hence mixed MAGUK oligomers may form via PDZ1-2 interactions, in the case of SAP97 with fewer restrictions on orientation due to membrane localization. Figure 6(a) shows the P2_1_3 packing with the sequence variation across PSD-95, SAP97, PSD-93 and SAP102 mapped onto the structure. It is evident that the PDZ1-2 scaffold could accommodate heterogeneity in the PDZ1-2 domains.

**Figure 6.**
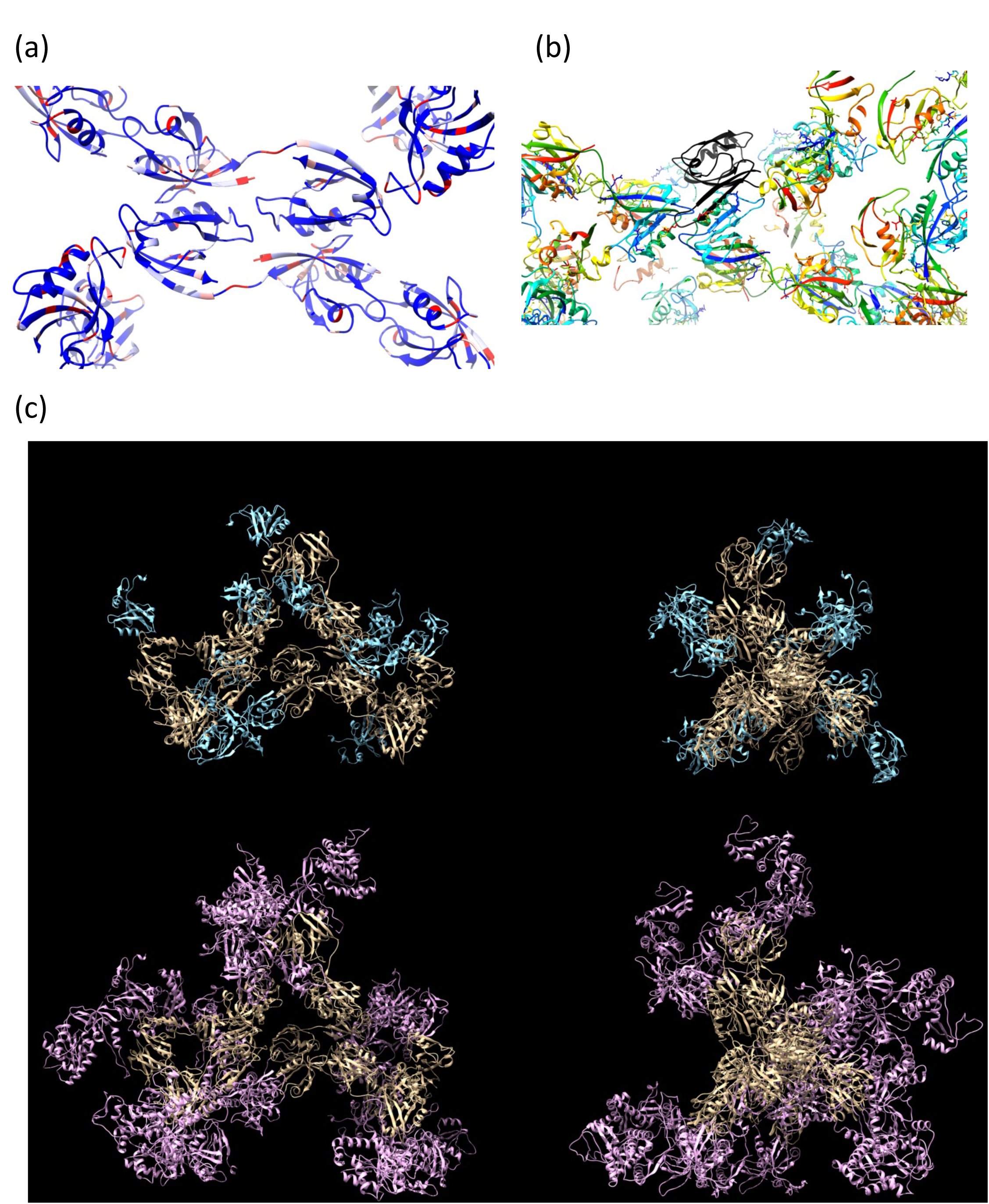
Additions to the PDZ1-2 scaffold. Sequence variation in the PDZ1-2 fragment across the MAGUK proteins is represented in (a). A sequence alignment obtained from the program ClustalW (Larkin et al, 2007) is mapped onto the packing arrangement with the locations of identical residues coloured blue, similar residues white and residues with different properties red. (b) shows the packing arrangement with PDZ1-2 displayed in a similar manner to Figure 4(a) and the nNOS PDZ domain (black) in association with PDZ1. The panels in (c) show two views of a dodecamer PDZ1-2-3 model and a PSD-95 model, the core PDZ1-2 scaffold is shown in yellow with PDZ3 in blue for the PDZ1-2-3 model and PDZ3-SH3GK in magenta for the PSD-95 model. All panels were generated with UCSF Chimera

Additional compatible PDZ domains can be incorporated into the PDZ1-2 scaffold replacing PDZ1 or PDZ2 in the scaffold Spacegroup. Figure 6(c) shows this process for the PDZ3 domain from PSD-95 (PDB ID 1bef: (Doyle et al, 1996)). PDZ3 can enhance the binding between tetramers by forming additional contacts to itself or to copies of PDZ1-2. In the model shown in Figure 6(c) this is the case for 8/12 PDZ3 domains in the modelled dodecamer, the PDZ3:PDZ1-2 contacts could be inter or intra-molecular in nature as restricted by the PDZ1-2-PDZ3 linker. Extending this formalism it is possible to gain some insight into the formation of whole-MAGuK oligomers. Using ZO-1 PDZ3-SH3-GK structure (Nomme et al, 2011) as a template, a model of the same region of PSD-95 can be realized by docking the structures of PDZ3 and SH3-GK (McGee et al, 2001). The modelled PDZ1-SH3-GK structure can then be docked onto the scaffold in the same way as for PDZ3. Athough as there is a clear conformational difference between the ZO-1 SH3-GK and PSD-95 SH3-GK any resulting analysis can only be approximate. This model is shown in Figure 6(c) using a (RRESEI:PDZ1):(RRESEI:PDZ3) contact for docking the PDZ1-SH3-GK addition. It is clear that the full-length protein can be incorporated into extended oligomers such as those observed in earlier Electron Microscopy images coincident with a 3_1_ screw axis. The linking of these together to form a net would require alternate conformations of PSD-95 or enzymatic cleavage to remove SH3-GK, or PDZ3-SH3-GK.

PSD-95 and SAP97 are substrates for the calpain family of calcium dependent proteases (Jourdi et al, 2005; Lu et al, 2000) which are implicated in synaptic plasticity (Baudry & Bi, 2016). Calpain cleavage of PSD-95 initially gives two fragments of PSD-95 of molecular weight ∼50 and 36kDa as measured by SDS-PAGE and western blotting. The sequence weights of these fragments and the epitope site of the antibody used in western blotting is consistent with PSD-95 cleavage in the linkers between PDZ2 and PDZ3 (liberating the ∼50kDa fragment) and between PDZ3 and SH3 (liberating the 36kDa fragment). The calpain-2 (catalytic) domain of the enzyme has a type I PDZ binding motif at the C-terminus (-FSVL) indicating that the enzyme can bind to type I PDZ domains to enhance the efficiency of target protein cleavage. Thus cleavage of MAGUKs by calpain may represent a calcium-dependent maturation process whereby the remnant PDZ1-2 can form oligomeric arrays. PSD-95 can be removed from the synapse via ubiquitination of a P/E/S/T rich motif: This sequence is located in the N-terminal linker region between the Cys-palmitoylation sites and the PDZ1 domain (Colledge et al, 2003). This indicates that a mechanism of removal for both full length and truncated PSD-95 in place, alongside a calpain degradation pathway together allowing a dynamic scaffolding structure to exist.

The core of a MAGuK scaffold layer can be formed by the association of ligated PDZ1-2 domains along directions parallel to two of the body diagonals of the scaffold unit cell as seen in Figure 5. Additional partner proteins may then integrate with this scaffold at available sites. The integration may be either via the incorporation of a compatible C-terminal sequence, or through the association of compatible PDZ domain(s) in the manner discussed above. A laminar model of the PSD (Valtschanoff & Weinberg, 2001) indicates that the nNOS protein is found in abundance at the cytoplasmic side of a PSD-95 enriched layer. The nNOS protein has an N-terminal PDZ domain that can afford integration with the PSD-95 scaffold. The PDZ domain of nNOS interacts with other PDZ domains via the insertion of a β-hairpin into the binding cleft as resolved in the crystal structure of the domain with syntrophin PDZ (Hillier et al, 1999). The equivalent interaction of the nNOS PDZ domain with the PDZ1 domain of PDZ1-2 in the scaffolding Spacegroup is shown in Figure 6(b). The binding cleft of PDZ1 is accessible in the PDZ1-2 scaffold and the β-hairpin can interact through association with the cleft. The PDZ1 domain is located at the rim of the voids shown in Figure 5(a), hence an association can form between nNOS and a core scaffold.

Modulation of the scaffold structure and in turn the underlying organization of channels may be afforded by relatively minor modifications to the ligand or the PDZ domain though post-translational modification. For instance phosphorylation of S/T residues in the type I PDZ sequence motifs are likely to affect the formation of oligomers. The Ser residue in RRESEI has a very high phosphorylation likelihood, as does the equivalent residue at the termini of the NMDA receptor (NetPhos 3.1 sever scores of 0.99 and 0.98 respectively (Blom et al, 1999)), and whilst a sequence bound at PDZ2 in the scaffold may be protected from kinases, one bound at PDZ1 is accessible as demonstrated for the addition of the nNOS PDZ domain.

## Conclusions

The PDZ1-2 component of PSD-95 is known to be an essential component for receptor clustering at the synapse and presumably other locations such as juxta-paranodes. This study shows that PDZ1-2 oligomerization can be accomplished by the dual domain and a ligating peptide alone – whole receptor proteins are not required. The PDZ domain itself may be considered to be an interacting element, interactions between PDZ domains are modulated by the properties of a bound peptide ligand. On binding a suitable peptide ligand particular structural elements of the PDZ1-2 domain are stabilized with mediation by the GLGF motif. and in turn favour the formation of oligomers.

The mechanism of clustering of PDZ1-2 is the formation of inter-molecular oligomers which follow a cubic packing rule and hence a highly organized scaffold underlies the formation of clusters. The ubiquitous intracellular peptide GSH can also induce the formation of inter-PDZ1-2 oligomers at concentrations comparable to those found in the cytoplasm. GSH may therefore be important in maintaining the fidelity of MAGUK induced clusters.

## Materials and Methods

### Protein production

All chemicals were obtained from Sigma-Aldrich (Fancy Road, Poole, Dorset BH12 4QH, UK) unless stated.

The sequence for PSD-95 PDZ1-2 (corresponding to residues 55-249 [98-292] of the UniProt ID P78352 [P78352-2]); PDZ1 55-152 [98-195] and PDZ2 154-249 [197-292] and PDZ3 303-415 [346-458], were cloned into a pOPINF expression vector (Berrow et al, 2007). The resulting protein is expressed with a HRV 3C protease cleavable hexa-histidine tag appended at the N-terminus of the protein. Tag-cleaved proteins have Gly-Pro followed by the protein sequence of interest. The sequence Molecular weight of the PDZ1-2 cleaved construct is 20.8 kDa.

Proteins were expressed after transformation into *E. coli* strain BL21(DE3) (New England Biolabs,75-77 Knowl Piece, Wilbury Way, Hitchin, Herts SG4 0TY UK.). Cultures were initially propagated in Double Yeast-Tryptone media supplemented with 50µg/ml Ampicillin at 37°C in shake flasks with 200rpm orbital shaking. Protein expression was induced at a culture optical density at 600nm of 0.6-0.8 by supplementing with 0.1mM isopropyl-β-D-thiogalactopyranoside. The cultures were then incubated at 16°C for 16-20 hours before harvesting of cells by centrifugation at 6,000×*g*. Cell pellets were flash frozen and stored at -70 °C.

Chromatography steps were carried out using an Akta FPLC system, controlled by UNICORN 5.01 software (GE Healthcare, The Grove Centre, White Lion Road, Amersham, Bucks, HP7 9LL, UK.). *E. coli* cells were resuspended in lysis buffer: 20 mM Na_2_HPO_4_/NaOH, 0.5 M NaCl, pH8.5, 1 mM reduced GSH, “Complete” mini EDTA-free protease inhibitor cocktail used at the manufacturer’s (Roche Diagnostics, Charles Avenue, Burgess Hill, West Sussex, RH15 9RY, UK.) recommended concentration, supplemented with 100 mg/l DNAse1. The cell suspension was sonicated (Bandelin Sonopuls HD3200, with TT13/F2 probe, Bandelin Electronic, GmbH & Co. KG, Heinrichstrasse 3-4, 12207 Berlin, Germany) on ice until the suspension was homogeneous. The lysate was then centrifuged at 39,000*g* to remove unbroken cells before application of the supernatant to a Ni-NTA agarose resin column (QIAGEN Ltd. Skelton House, Lloyd St. North, Manchester, M15 6SH, UK). The affinity column was washed with lysis buffer supplemented with 10mM imidazole to remove weakly interacting proteins. Immobilized protein was then eluted in a stepwise manner with lysis buffer supplemented with 200mM imidazole. Desalting chromatography was performed using a HiPrep 26/10 column (GE Healthcare). Tag cleavage was carried out overnight at 4 °C on a roller with 10 units of HRV 3C protease (Novagen/Merck chemicals Boulevard Industrial Park, Padge Road, Beeston, Nottingham, NG9 2JR, UK) per mg of protein. A negative Ni-NTA affinity purification step was carried out to remove uncleaved product, the flow through from this step was concentrated using a centrifugal concentrator and applied to a Superdex200 10/300 GL (GE Healthcare) size exclusion column equilibrated with a “Standard” buffer of 20 mM Tris-HCl, 150 mM NaCl, pH8.5. PDZ1-2, PDZ1 and PDZ2 showed a single band on a Coomassie stained SDS-PAGE gel and a single highly symmetric peak in size exclusion at a retention volume consistent with the expected Molecular Weight of the construct.

### X-ray crystallography

#### Crystallization and Data collection

PDZ1-2 protein from a single size exclusion chromatography peak was pooled and concentrated with centrifugal concentrator to a concentration of ∼0.75mM. Vapour diffusion sitting drop 96-well crystallisation plates were set up using an automated liquid handler (Mosquito Crystal, ttp Labtech, Melbourn Science Park, Melbourn, Herts, SG8 6EE, UK.). The volume ratio of protein and screen solution was 200:200nL. Each drop was equilibrated against a 100µL well volume at 4 °C. Crystals of Apo:PDZ1-2 with a maximum dimension of 250 µm and a tetragonal bipyramid habit were observed after 4 weeks from crystallization with a screen solution containing 0.2M calcium acetate, 0.1M sodium cacodylate pH6.5, 40% v/v PEG300.

A number attempts were made to crystallize RRESEI:PDZ1-2 without a seeding step, these were unsuccessful. A large number of clear wells were seen in crystal screens, which is consistent with high solubility of the protein-ligand complex limiting nucleation. Start and ending protein concentrations in the vapour diffusion based crystallization trials were ≈0.5 and 1 mM respectively. PDZ1-2 crystals in the presence of ligand were obtained via Matrix micro-seeding (D’Arcy et al, 2007). A PDZ1-2 solution containing an excess concentration of the ligand peptide RRESEI at 98% purity (Peptide Protein Research Ltd., Bridge House Farm, 184 Funtley Road, Funtley, Fareham PO15 6DP, UK.) was prepared by supplementing a 1mM solution of PDZ1-2 with 10mM of the RRESEI peptide and incubating for 4 hours at 4 °C on a roller. Crystallisation trials were set up using a Mosquito (TTP Labtech) liquid handling robot, with screens obtained from Molecular Dimensions (Unit 6 Goodwin Business Park, Willie Snaith Road, Newmarket, Suffolk, CB8 7SQ, UK.). The seed stock was prepared by mixing a drop containing Apo:PDZ1-2 **c**rystals (volume ≤ 400 nl) with 350 μl of the corresponding reservoir and pulverizing the crystal using a micro-seed bead (Molecular Dimensions). The seed stock, the protein, and the crystallisation screen reagent were dispensed consecutively to form drops comprising 150 nl protein solution: 50nl seed suspension: 200 nl reservoir. Crystals of maximum dimension 250 µm with a tetragonal bipyramid habit were observed after 1 week in two screen solutions containing (a) 0.2 M NaCl, 0.1 M Na/K phosphate, pH 6.2, 50% v/v PEG200; and (b) 0.2M Li_2_SO_4_, 0.1M TRIS pH8.5, 40% v/v PEG400.

For both Apo:PDZ1-2 and PDZ1-2+RRESEI crystals were harvested into fibre loops and flash cooled in liquid nitrogen directly from crystallization drops. Diffraction data were collected from a single crystal at the Diamond Light Source. Crystallographic data was processed using the Xia2 expert system, CCP4 software and XDS integration software (Collaborative Computational Project, 1994; Evans, 2006; Kabsch, 2010; Winter, 2010; Winter et al, 2013). Subsequent data analysis was facilitated by programs from the CCP4 suite. The crystal structure reported here for PDZ1-2+RRESEI crystals is from the PEG200 condition given above which showed higher resolution diffraction (2.1 versus 2.4 Å).

#### Crystallographic structure solution, model building and refinement

Crystal structures of PDZ1 and PDZ2 from human SAP-97 (Zhang et al, 2011)(PDB IDs: 3rl7, 3rl8) have been determined at high resolution in complex with a ligand peptide with sequence RHSGSYLVTSV. The two PDZ domains have a high level of sequence identity with their counterparts in PSD-95. Molecular replacement was carried out with the models prepared from one domain from each of these structures, using the Chainsaw program (Stein, 2008) and the sequence of the expressed construct. Molecular replacement of the Apo crystal structure was effected with the program PHASER (Mccoy et al, 2007) and gave a solution with Log-likelihood gain of 771 and one copy each of PDZ1 and PDZ2 in the asymmetric unit. Clear electron density for unique residues in the individual domains was observed as was connecting density for the linker region between the two domains. The resulting initial model was examined to identify the linked PDZ1-2 double domain. The Ligand soaked crystal structure was solved by cross phasing using a partially refined Apo:PDZ1-2 model, followed by rigid body refinement. For the crystals grown in the presence of the RRESEI ligand, clear and unbroken electron density was seen for the main chain of ESEI residues in the initial Fo-Fc difference electron density map. Model building was carried out using the program Coot (Emsley et al, 2010), interspersed with refinement using REFMAC5 (Murshudov et al, 1997). In the refinement process each of the domains in PDZ1-2 was assigned to a rigid body and tensors describing Translation, Libration and their correlation were used in REFMAC5 to describe anisotropy in the model (Winn et al, 2001). Data collection and refinement parameters are included in the Supplementary Table 1.

#### Isothermal Titration Calorimetry

Isothermal Titration Calorimetry (ITC) experiments were carried out using a MicroCal VP-ITC (Malvern/Panalytical Ltd., Enigma Business Park, Grovewood Road, Malvern, WR14 1XZ, UK.) instrument. All samples were prepared in the aforementioned Standard buffer and this buffer was used in control experiments to measure the heat change on dilution. A 5mM concentration of the RRESEI ligand was placed in the syringe with either PDZ1-2; PDZ1 or PDZ2 placed in the cell at a nominal concentration of 0.13mM. ITC experiments were conducted at 25°C. After an initial priming injection, 19 consecutive 10µL aliquots were injected into the cell each over 4.8s and the heat change recorded over a 250s interval before the next injection. Data analysis was performed using OriginR 5.0 software (OriginLab, Silverdale Scientific Ltd., Silverdale House, 111 Wendover Road, Stoke Mandeville, Bucks, HP22 5TD, UK.).

### Small Angle X-ray Scattering

#### SAXS data collection

Small Angle X-ray Scattering (SAXS) data was collected on PDZ1-2 in two modes at two beamlines. In the case where the RRESEI peptide ligand was present the protein sample was prepared in the same way described for protein crystallization. In the cases where reduced Glutathione (GSH) is present the protein sample was diluted with a 20mM concentration of GSH prepared in Standard buffer to achieve the stated concentrations. Unfractionated SAXS data was collected on 20µL solution samples at fixed concentrations alongside the corresponding buffer reserved from the final chromatography step. For Size Exclusion Chromatography SAXS (SEC-SAXS), scattering experiments were carried out after fractionation using an Agilent 1260C HPLC system (Agilent Technologies LDA UK Ltd., Life Sciences and Chemical Analysis group, Lakeside, Cheadle Royal Business Park, Stockport Cheshire, SK8 3GR, UK.), and data were collected on samples eluted from a 4.6ml Shodex Kw403 silica chromatography column (Showa Denko Europe GmbH, Konrad-Zuse-Platz 3, 81829 Munich, Germany). Two collection modes were used for SEC-SAXS: (1) where the liquid chromatography flow is paused and data images are recorded and (2) where data is continuously recorded and images selected corresponding to SEC peak fractions are subsequently combined. In the SEC-SAXS case the buffer data for background subtraction was selected from frames within the size exclusion run.

Data images were recorded on area detectors and reduced to 1-dimensional scattering profiles by beamline software. These scattering profiles were compared within each data set and those showing evidence of radiation damage (diagnosed by enhanced scattering at low q at the expense of high q scattering (Jacques et al, 2012)) excluded before the profiles were merged using PRIMUS (Konarev et al, 2003). Supplementary Table 2 summarizes the SAXS data sets recorded. Detailed results are shown for the unfractionated SAXS at highest resolution for Apo:PDZ1-2 and RRESEI:PDZ1-2 and GSH:PDZ1-2, and for SEC-SAXS SAXS with paused chromatography for Apo:PDZ1-2 and RRESEI:PDZ1-2 and GSH:PDZ1-2. Equivalent results were found for the integrated peak SAXS (Jun ’16) and lower resolution unfractionated native (Sept’16).

#### Initial analysis of SAXS profiles

Data were collected on Apo:PDZ1-2, RRESEI:PDZ1-2 and GSH:PDZ1-2. In each case data quality was excellent with low noise levels even for data collected on diluted samples at higher q values. The scattering profiles obtained were significantly different according to exposure to RRESEI ligand or the presence of GSH. Initial data analysis showed that for the unfractionated SAXS data in all cases the Guinier regions were non-linear. The estimates of maximum pair distance (D_max_) values obtained from the Pair distance distribution function also differed with the concentration of the sample, with larger values of D_max_ correlating with higher concentrations of PDZ1-2. In each case (Apo/RRESEI/GSH) the fractionated (SEC-)SAXS data indicated a more homogenous sample when compared to the unfractionated SAXS data.

The scattering curves recorded in the unfractionated SAXS case indicated that inter-particle interaction was occurring (Jacques & Trewhella, 2010). Comparing the scaled scattering curves at different concentrations at low values of q indicated that a very small amount of “repulsive” inter-particle interference was present in the Apo:PDZ1-2 case (a lower I(q) compared to the curve recorded after dilution). A much larger “attractive” interference (a raised I(q) compared to the curve recorded after dilution) was observed for RRESEI:PDZ1-2. There are no Cys residues in the sequence of the PDZ1-2 expressed protein hence the direct association of PDZ1-2 via disulphide bonding is not possible. In each case Kratky plots of the SAXS data did not indicate significant overall change in the folding of the protein (Rambo & Tainer, 2011). In all cases the symptoms of inter-particle interactions were alleviated by dilution, therefore the inter-particle interactions observed are reversible. The inter-particle interactions are consistent with non-covalent associations between copies of PDZ1-2. Initial analysis was performed using unfractionated SAXS data collected on Apo:PDZ1-2 and RRESEI:PDZ1-2. The data at fixed concentrations (≈0.72 and 0.35mM) were projected to zero concentration using PRIMUS (Konarev et al, 2003) to minimize the inter-particle effects on the scattering curves (Figure 7(a,b)).

**Figure 7.**
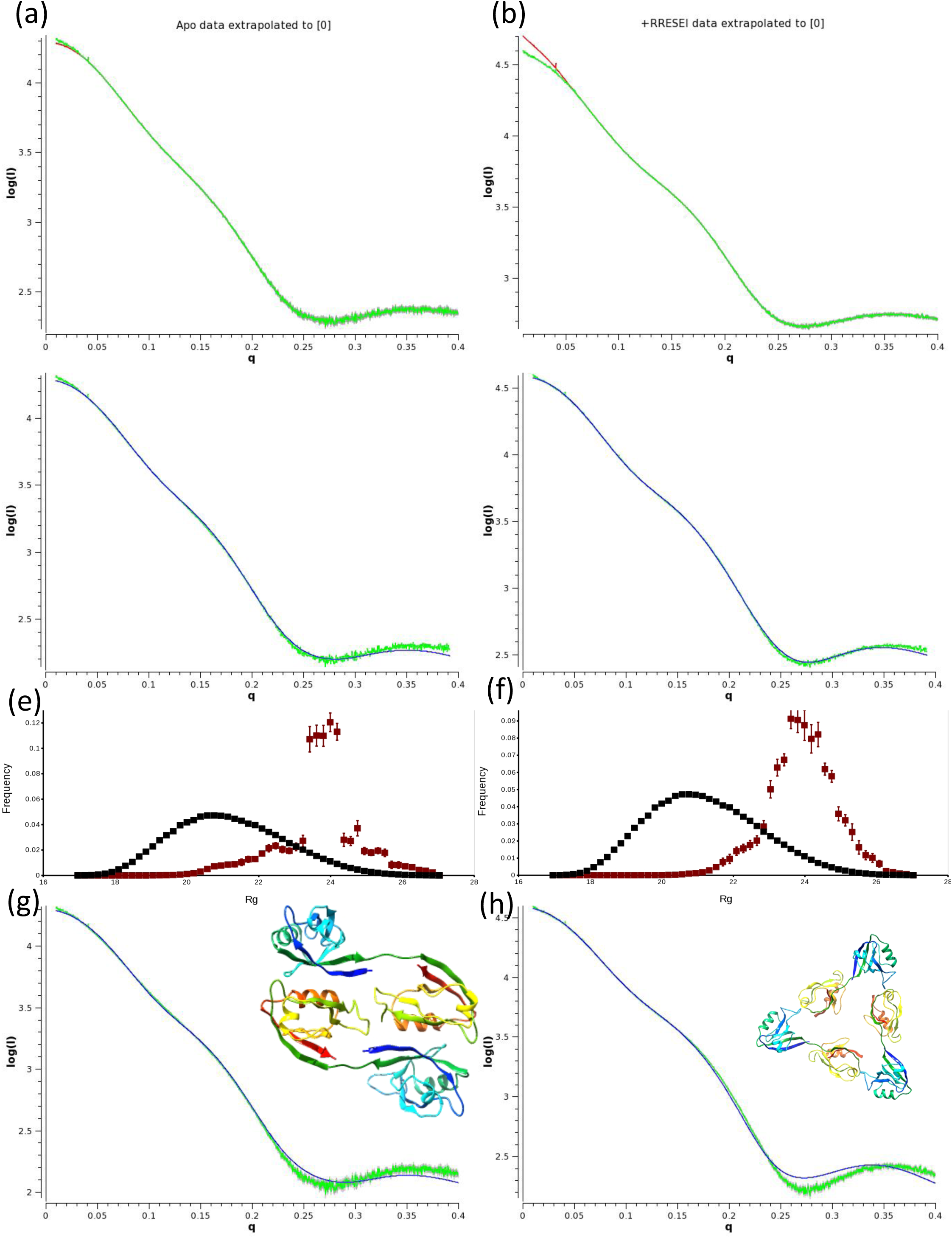
Initial Model based analysis of apo:PDZ1-2 and RRESEI:PDZ1-2 SAXS data. Scattering data are plotted as Log(I) versus momentum transfer with the projected data in green and calculated fitted curves shown in blue. The extrapolation of data to infinite dilution for apo:PDZ1-2 and RRESEI:PDZ1-2 recorded at the DESY synchrotron in June ’15 (see supplementary Table 2) are shown in (a) and (b) respectively, with high concentration data shown in red. (c) and (d) show the Ensemble model analysis fit (Bernado et al, 2007), with the selected model histogram in (e) and (f): Population is plotted on the ordinate and R_g_ on the abscissa, black points are pool models and red points models selected from the pool, with error bars extracted from 20 duplicate runs. (g) and (h) show monomer-oligomer refinements (Petoukhov et al, 2012) in each case an extended monomer (Figure 4(a)1e) is refined alongside the oligomer shown as an inset.

Analysis using dummy atom models (DAMMIF) (Franke & Svergun, 2009) was undertaken The agreement with the data curves reasonable although data were truncated (q ≤ observed R_g_/8) by DAMMIF to allow dummy atom modelling. For data collected in the presence of RRSEI the data range was 0.015 < q < 0.300 Å^-1^, Χ^2^ value 8.7; in the Apo case 0.011 < q < 0.204 Å^-1^, Χ^2^ = 6.7. The modelling gave envelopes which would accommodate the PDZ1-2 model but with additional dummy atom distributions (Supplementary Figure 1).

#### Model based analysis of SAXS data

Model based analysis with the models from Ensemble Optimization Method (EOM) (Bernado et al, 2007) software using the data extrapolated to zero, was carried out with a limit of q=0.4Å. The analysis gave different histogram distributions of model Radius of gyration according to the presence or absence of RRESEI: A sharper R_g_ histogram distribution for Apo:PDZ1-2 data compared to a broader population for RRESEI:PDZ1-2 (Figure 7(e,f)). The agreement of the models derived from the EOM analysis was best over the mid-q range of the curve and poorer at higher and low q values (Figure 7(c,d)). The representative models generated by EOM generally contained a model with an R_g_ of around 25Å with a high weight assigned, alongside one or more compact models (R_g_ in the range 21-24Å).

EOM analysis at fixed concentrations (≈0.72 and 0.35mM) for the Apo:PDZ1-2 gave good agreement over the mid-range of data (Χ^2^ values for the concentrated data ∼5). However the EOM histograms gave bifurcated distributions of models with different weights of the two peaks for different concentrations. The minor peak at R_g_ 22.5Å shown for the Apo extrapolated data histogram (Figure 7(e)) is increased at the expense of the major peak around 24Å. In the case of the RRESEI:PDZ1-2 at fixed concentrations (≈0.72 and 0.35mM) the agreement obtained with the data by EOM was very poor (Χ^2^ values for the 0.72mM data > 999), with large differences at both low and high q. The model histograms gave a single broad peak at each concentration, but the peak was skewed toward higher R_g_ values for the data at higher concentration.

#### Construction of Oligomer models for SAXS fitting

Oligomer models were constructed from combinations of a known conformation of PDZ1-2 and known inter-molecular interfaces derived from X-ray crystal structures. All of the PDZ1 and PDZ2 domains in both the 3gsl and 3zrt crystal structures take part in the same PDZ1:PDZ2 heterodimeric interaction shown in Supplementary Figure S2(a). The αB helix adjacent to the unoccupied peptide binding cleft of each PDZ2 domain associates with the βD-βE loop from PDZ1. The significance of this interaction is shown by the fact that the 3zrt structure can be solved by molecular replacement using an ensemble made from the two PDZ1:PDZ2 heterodimers present in the asymmetric unit of 3gsl: Using the CCP4 program PHASER (Mccoy et al, 2007) and the deposited Crystallographic data a Log Likelihood gain of 1337 was obtained for 4 copies of the ensemble: The Log Likelihood gain value may be enhanced by the Non-crystallographic symmetry implied by the presence of multiple copies of the heterodimer, but it is much greater than the threshold of 8 which indicates a correct solution. Therefore this αB(PDZ2):βD-βE(PDZ1) interface was used as an oligomeric contact. One obvious oligomer is the double dimer of which there are 2 copies in the asymmetric unit of 3zrt:PDZ1-2.

Both PDZ1-2 crystal structures reported were found to have an *<u>intra</u>*-molecular contact formed between the βB-βC loop of PDZ1 and the αA helix of PDZ2. A similar *<u>inter</u>*-molecular contact was found in the 3gsl structure with the αA helix of PDZ2 forming a contact with the βB-βC loop of PDZ1 (in 3gsl a Cα-Cα distance of 5.3Å is obtained between Ala 199 in chain A and Asp 90 of chain B after the application of the symmetry operation 1-x, ½+y,-z). This αA(PDZ2): βB-βC(PDZ1) crystal contact (shown in Supplementary Figure S2(b)), was also used as an oligomeric contact. On combining the αA(PDZ2): βB-βC(PDZ1) interaction with the 3zrt-like conformation of PDZ1-2 it was clear that a trimeric complex could also form.

Initial modelling of apo:PDZ1-2 and RRESEI:PDZ1-2 SAXS curves projected to zero concentration was undertaken using these models. Monomer/oligomer fractions were refined using the ATSAS program SASREFMX (Petoukhov et al, 2012) and encouraging results were obtained for both Apo:PDZ1-2 and RRESEI:PDZ1-2 curves projected to zero concentration (Figure 7(g,h)). In the Apo:PDZ1-2 case the combination of 3zrt:PDZ1-2 along with a dimer of the same (similar to one half of the asymmetric unit of the 3zrt:PDZ1-2 crystal structure, Figure 7(g) inset) was effective in SAXS curve fitting at low-medium resolution (Figure 7(g)) and gave Χ^2^ values of ≈16. In the case of RRESEI:PDZ1-2 the 3zrt:PDZ1-2 conformation plus a trimer of the same (Figure 7(h) inset) was similarly effective (Χ^2^ values of ≈28, Figure 7(h)). The SASREFMX analysis also provided a refined model for a trimer of RRESEI:PDZ1-2. The application of this simple oligomer model was also more effective in fitting the RRESEI:PDZ1-2 curves at fixed concentrations giving a marked improvement in agreement (Χ^2^ values ≈60 and 30 for 0.72 and 0.36mM scattering curves respectively) over the previous EOM based analysis. Using the dimer interface seen in 3zrt:PDZ1-2 along with the trimer determined by SAXREFMX allows the generation of extended oligomers. It was found that a trimeric oligomer of extended PDZ1-2 could interact with a copy of itself through the formation of a dimer-of-trimers, with a local interface equivalent to the double dimer found in the 3zrt structure. These types of interactions could then be propagated to form more extended oligomers including the formation of a trimer of trimers with 3-fold screw-rotational symmetry. Further combinations of these trimers indicated that extended arrays could be formed, including linear and branched arrays. The inclusion of these types of oligomers in SAXS fitting of RRESEI + PDZ1-2 data at fixed concentrations using the ATSAS program OLIGOMER (Franke et al, 2017; Konarev et al, 2003) again improved the fit of the data.

#### Assignment of the I2_1_3 Spacegroup to the packing arrangement of the PDZ1-2 oligomers

Exploring the structures formed by the association of trimers of PDZ1-2 revealed the presence of parallel 3-fold rotation and 3_1_ screw axes and, separately, parallel 2-fold rotation axes each in orthogonal directions. These symmetry elements are present in multiple directions only in cubic Space Groups (Numbers 195 to 230 in International Tables for Crystallography volume A (Hahn, 1983)). From the set of 13 chiral cubic Space Groups those possessing 4 or 4_1_ axes could be eliminated, and as the 3-fold axes do not intersect in the extended oligomers, and 2-fold axes are present the only possible Spacegroup conforming to the symmetry elements encountered is I2_1_3 (Number 199) (Hahn, 1983).

The unique unit cell parameter (<u>a</u>) for the cubic lattice and the position of the PDZ1-2 domain with respect to the coordinate origin then required definition. This was accomplished via generating a trimeric arrangement of 3 PDZ1-2 double domains. Firstly a model of the PDZ1-2 double dimer was obtained. Domains from the PDZ1-2:RRESEI crystal structure were docked onto the 3zrt:PDZ1-2 conformation and restrained refinement of this structure against RRESEI:PDZ1-2 SEC-SAXS data was carried out using SASREFMX. Inter-domain restraints were obtained from the αB(PDZ2):βD-βE(PDZ1) interface in the 3gls:PDZ1-2 crystal structure and monomer + symmetric dimer model was refined. The resulting dimer was similar in form to those found in the 3zrt crystal structure but with a slightly increased twist of the dimer about the interface perpendicular to the 2-fold axis. The twofold rotational symmetry axis of this dimer was then oriented with reference to a set of mutually perpendicular right-handed axes x, y and z such that, the 2-fold axis was in a direction perpendicular to the z/y plane and intersecting the x axis. A trimer of dimers can then be generated by rotation about the resultant vector of the x, y and z axes. The rotation of the dimer about the 2-fold axis and the displacement of the dimer from the z/y plane Δx, were adjusted so that the contact between dimers in the trimer was similar to the αA(PDZ2): βB-βC(PDZ1) crystal contact found in 3gsl:PDZ1-2. The position of PDZ1-2 was then fixed as one copy within the dimer of trimers and |**<u>a</u>**| the unit cell length could then be defined as 4×(Δx)=148Å. The precision of this process is dependent upon the accuracy of the dimer model and the positioning of this model in space. When generating the trimer of dimers the interval of rotation about the 2-fold axis was 5° and the interval in Δx was 1Å. Thus the imprecision is likely to be dominated by the error in the SASREFMX refinement of the dimer at a q(max) of 0.4Å^-1^ - on this basis the error is estimated to be (2π)/(2×q(max)) ≈ 8Å.

Using the I2_1_3 Spacegroup with cell length |**<u>a</u>**|=148 Å, alongside the origin determined as described above allowed any packing arrangement to be faithfully reproduced based on a single copy of the extended PDZ1-2 molecule. The two inter-molecular contacts namely the αB(PDZ2):βD-βE(PDZ1) and the αA(PDZ2): βB-βC(PDZ1) are features of this lattice (Supplementary Figure 2(a,b)). For the final model the αB(PDZ2):βD-βE(PDZ1) PDZ1:PDZ2 dimer was re-docked into the unit cell in order to preserve the fidelity of the interface precisely. Minor side-chain clashes in models were resolved by adjusting rotamers in COOT followed by geometrical refinement in REFMAC5.

## Acknowledgements

X-ray crystallographic data was collected at Diamond Light Source Beamlines I04 and I04-1. ITC was conducted in the Biophysical analysis facility of the University of Manchester. SAXS data was collected at the DESY Synchrotron Beamline P21 and at Diamond Light Source Beamline B21. NAR was supported by a Majlis Amanah Rakyat Malaysian Government Scholarship. JC was supported by a BBSRC/EPSRC Doctoral Training award. CB and MLC are members of the Wellcome Trust Centre for Cell-Matrix Research, supported by funding from the Wellcome Trust (Ref. 203128/Z/16/Z). CB gratefully acknowledges BBSRC funding (Refs: BB/N015398/1 and BB/R008221/1).

## Author contributions

LB designed and generated the clones and characterized their expression in *E.coli* bacteria at the Oxford Protein Production Facility and with support from the Oxford Module Consortium; JC corrected cloning artefacts via site directed mutation with advice from Dr. James Birtley (University of Manchester) and carried out pilot purification with the help of Dr. Ronald Burke (Manchester Protein Production facility). NAR produced and purified the protein samples for each of the studies reported here. Protein crystallization was performed by NAR and CWL; X-ray Crystallographic data was collected by CWL. Crystallographic interpretation was carried out by NAR & SMP. ITC was conducted by NAR. CB and MLC collected SAXS data and carried out initial interpretation. SAXS interpretation was carried out by NAR & SMP. The information provided by the 3gls and 3zrt structures of PDZ1-2 was essential to the analysis of SAXS data and the resulting oligomer models presented here. The manuscript was written by SMP with contributions from NAR, CB and CWL.

## Data availability

The Apo:PDZ1-2 and RRESEI:PDZ1-2 refined structures have been deposited in the protein data bank (https://www.wwpdb.org) with PDB accession codes 6spv and 6spz. The concentrated, diluted and SEC-SAXS curve data with corresponding oligomer models have been deposited in the SASBDB (https://www.sasbdb.org): For RRESEI:PDZ1-2 accession codes SASDGB5, SASDGC5, SASDGD; Apo:PDZ1-2, SASDGE5, SASDGF5, SASDGG5 and GSH:PDZ1-2 SASDGH5, SASDGJ5, SASDGK5. A PDB format file representing the asymmetric unit of the PDZ1-2 scaffold can be generated from the chain identifier “A” of any of the oligomer models deposited in the SASBDB. The header can be generated using Spacegroup symmetry P2_1_3 with the unique cell parameter |**<u>a</u>**|=148 Å.

## Conflict of Interest

The authors declare no conflict of interest.

